# Chronic social defeat causes dysregulation of systemic glucose metabolism via the cerebellar fastigial nucleus

**DOI:** 10.1101/2025.02.18.638938

**Authors:** Taiga Ishimoto, Takashi Abe, Yuko Nakamura, Tomonori Tsuyama, Kunio Kondoh, Naoto Kajitani, Kaede Yoshida, Yuichi Takeuchi, Kan X. Kato, Shucheng Xu, Maru Koduki, Momoka Ichimura, Takito Itoi, Kenta Shimba, Yoshifumi Yamaguchi, Masabumi Minami, Shinsuke Koike, Kiyoto Kasai, Jessica J Ye, Minoru Takebayashi, Kazuya Yamagata, Chitoku Toda

**Affiliations:** Department of Neuroscience for Metabolic Control, Graduate School of Medical Science, Kumamoto University, Kumamoto, Japan; Center for Evolutionary Cognitive Sciences, Graduate School of Art and Sciences, The University of Tokyo, Tokyo, Japan; University of Tokyo Institute for Diversity & Adaptation of Human Mind (UTIDAHM), Tokyo, Japan; Center for Metabolic Regulation of Healthy Aging (CMHA), Faculty of Life Sciences Kumamoto University, Kumamoto, Japan; Division of Endocrinology and Metabolism, Department of Homeostatic Regulation, National Institute for Physiological Sciences, National Institutes of Natural Sciences, Aichi, Japan; Division of Integrative Physiology, Faculty of Medicine, Tottori University, Tottori, Japan; Department of Neuropsychiatry, Faculty of Life Sciences, Kumamoto University, Kumamoto, Japan; Department of Pharmacology, Graduate School of Pharmaceutical Sciences, Hokkaido University, Hokkaido, Japan; Department of Pharmacy, Faculty of Pharmacy, Kindai University, Osaka, Japan; Laboratory of Biochemistry, Graduate School of Veterinary Medicine, Hokkaido University, Hokkaido, Japan; Graduate School of Frontier Sciences, The University of Tokyo, Chiba, Japan; Hibernation Metabolism, Physiology and Development Group, Institute of Low Temperature Science, Hokkaido University, Hokkaido, Japan; The International Research Center for Neurointelligence (WPI-IRCN), Institutes for Advanced Study (UTIAS), University of Tokyo, Tokyo, Japan; Department of Neuropsychiatry, Graduate School of Medicine, International Research Center for NeuroIntelligence, The University of Tokyo, Tokyo, Japan; Department of Neurology, Brigham and Women’s Hospital, Harvard Medical School, Boston, MA, USA; Department of Medical Biochemistry, Faculty of Life Sciences Kumamoto University Kumamoto, Kumamoto, Japan

## Abstract

Chronic psychological stress leads to hyperglycemia through the endocrine and sympathetic nervous systems, which contributes to the development of type II diabetes mellitus (T2DM). Higher plasma corticosteroids after stress is one well-established driver of insulin resistance in peripheral tissues. However, previous studies have indicated that only a fraction of patients with depression and post-traumatic disorder (PTSD) who develop T2DM exhibit hypocortisolism, so corticosteroids do not fully explain psychological stress-induced T2DM. Here, we find that chronic social defeat stress (CSDS) in mice enhances gluconeogenesis, which is accompanied by a decrease in plasma insulin, an increase in plasma catecholamines, and a drop in plasma corticosterone levels. We further reveal that these metabolic and endocrinological changes are mediated by the activation of neurons projecting from the cerebellar fastigial nucleus (FN) to the medullary parasolitary nucleus (PSol). These neurons are crucial in shifting the body’s primary energy source from glucose to lipids. Additionally, data from patients with depression reveal correlations between the presence of cerebellar abnormalities and both worsening depressive symptoms and elevated HbA1c levels. These findings highlight a previously unappreciated role of the cerebellum in metabolic regulation and its importance as a potential therapeutic target in depression, PTSD, and similar psychological disorders.

---

T2DM is one of the most common and costly conditions in the developed world and its incidence continues to increase worldwide^1^. While diet and exercise are commonly recognized to influence the development of T2DM, depression and PTSD have also been shown to be important risk factors^2–4^. In this study, we use a mouse model of chronic social defeat stress (CSDS) to investigate the neuro-endocrine mechanisms of stress-induced hyperglycemia. Mice subjected to CSDS do not exhibit glucose intolerance immediately after the 10-day stress period, but develop abnormalities about one week after the cessation of stress exposure^5^. As such, it is an ideal model to elucidate the mechanisms by which stress alters glucose metabolism in peripheral tissues.

## Glucose intolerance is induced in the post-CSDS period independently of the HPA axis

To induce CSDS, C57BL/6 mice were exposed to dominant ICR mice for 15 min/day, then separated with a wire mesh in the same cage, then re-exposed to ICR mice repeatedly in the same manner for a total of 10 days (Fig. 1a). As expected, mice developed glucose intolerance one week after CSDS exposure (referred to as the post-CSDS period), but not immediately after CSDS (referred to as the immediate-CSDS period) (Fig. 1b, c). Conversely, plasma corticosterone concentration increased immediately after CSDS, but decreased in the post-CSDS period when the mice started showing signs of glucose intolerance (Fig. 1d). CSDS-exposed mice also had increased avoidance behavior during social interaction (SI) testing (Extended Data Fig. 1a) and spent less time in the center during open field (OF) testing, both immediately after CSDS and during the post-CSDS period (Extended Data Fig. 1b, c). During the post-CSDS period, time spent in the interaction zone during SI testing, and time spent in the central zone during OF testing were also both negatively correlated with blood glucose levels during glucose tolerance testing (GTT), suggesting that the degree of glucose-induced hyperglycemia correlates with severity of anxiety behaviors (Extended Data Fig. 1d, e). Of note, however, this occurs only in the post-CSDS phase when corticosteroid levels have already dropped, suggesting it occurs independently of the HPA axis.

**Fig. 1.**
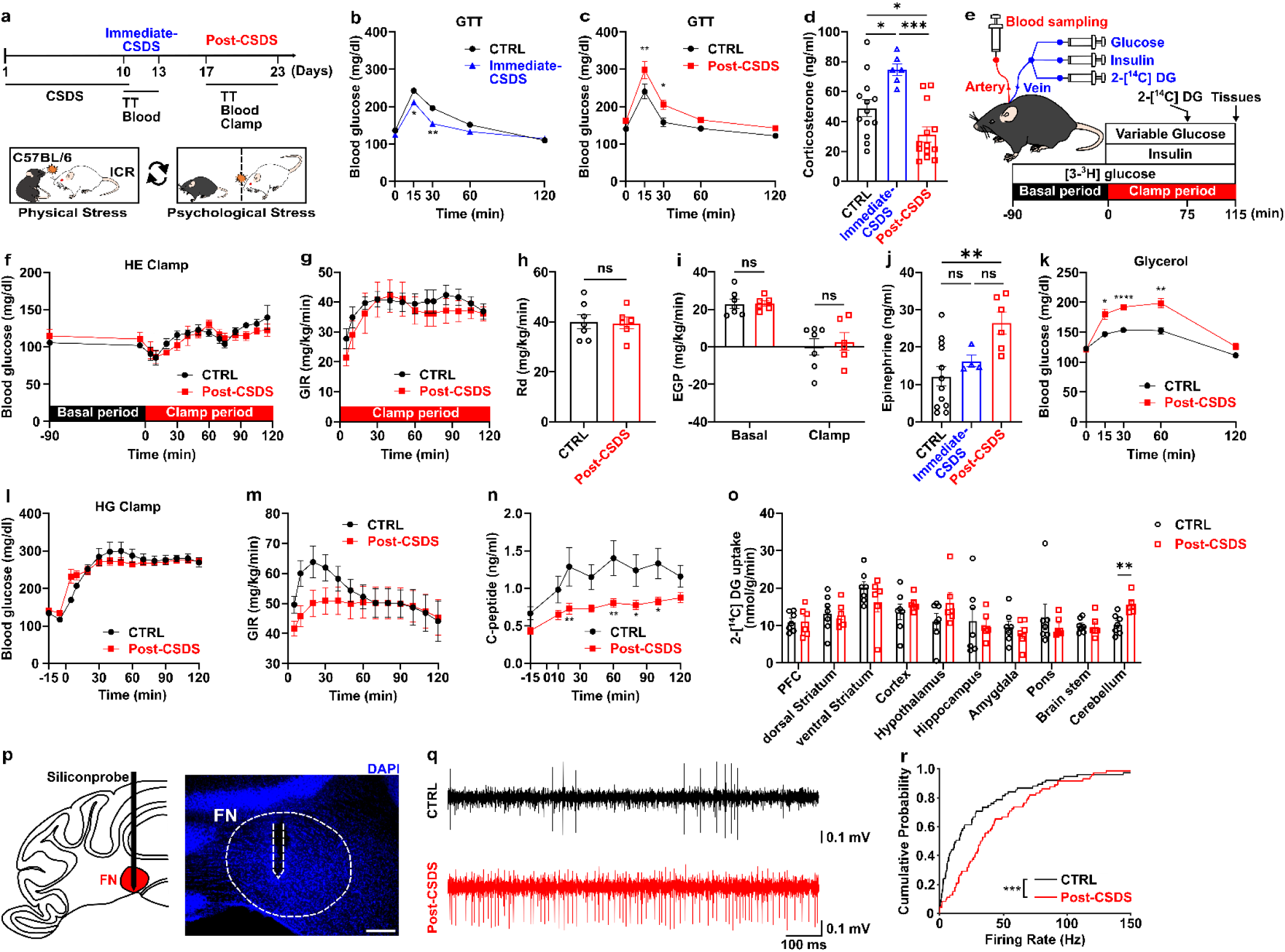
CSDS promotes gluconeogenesis by increasing plasma epinephrine during post-CSDS. **a**, Experimental timeline for the metabolic test. Chronic social defeat stress (CSDS) was exposed to C57BL/6 through a 10-day cycle. We defined days 10–13 as the immediate-CSDS period and days 17–23 as the post-CSDS period. TT, glucose or glycerol tolerance tests; Blood, blood sampling. **b**, Glucose tolerance test (GTT) of control (CTRL, n = 12) or immediate-CSDS (n = 14) mice. **c**, GTT of control (n = 8) or post-CSDS (n = 7) mice. **d**, Plasma corticosterone levels of control, immediate-CSDS, and post-CSDS mice (n = 13, 6, 13). **e**, schematic illustration of the hyperinsulinemic-euglycemic clamp (HE Clamp) procedure. The HE Clamp consists of the basal period and clamp period. 2-[^14^C] DG, 2-[^14^C]-Deoxy-D-Glucose. **f-i**, Blood glucose levels (**f**), glucose infusion rate (GIR, **g**), glucose disappearance (Rd, **h**), and endogenous glucose production (EGP, **i**) during HE Clamp (CTRL n = 7, post-CSDS n = 6). **j**, Plasma epinephrine levels of control, immediate-CSDS, and post-CSDS mice (n = 12, 4, 6). **k**, Glycerol tolerance test (0–120 min) of control (n = 9) or post-CSDS (n = 8) mice. **l-n**, Hyperglycemic clamp (HG Clamp) in control (n = 8) and post-CSDS (n = 16) mice. Blood glucose levels (**l**), GIR (**m**), plasma C-peptide levels (**n**, post-CSDS n = 15) during HG Clamp. **o**, 2-[^14^C] DG uptake of each brain region in HE Clamp. **p**, Representative position of inserted siliconprobe into the fastigial nucleus (FN). The scale bar is 200μm. **q**, Representative in vivo extracellular recordings in control and the post-CSDS mice. **r**, The Cumulative probability of firing rate in control (n = 75 neurons from six mice) and post-CSDS (n = 72 neurons from three mice) mice. Data are presented as mean ± SEM; \**p* < 0.05, \*\**p* < 0.01, \*\*\**p* < 0.001, \*\*\*\**p* < 0.0001, two-way ANOVA followed by Sidak multiple comparison tests in **b**, **c**, **f**, **g**, **k-n**; one-way ANOVA followed by Tukey’s multiple comparison tests in **d**, **i**, **j**; two-tailed t-test in **h**, **o**, two-sample Kolmogorov-Smirnov test in **r**.

A decrease in whole-body glucose utilization is often linked to reduced insulin sensitivity. However, the insulin tolerance test showed no changes in insulin sensitivity during the immediate- or post-CSDS period (Extended Data Fig. 2a, b). To further assess insulin sensitivity, we performed a hyperinsulinemic-euglycemic clamp (HE-Clamp) (Fig. 1e). In the post-CSDS period, whole-body glucose utilization remained unchanged. Glucose uptake in peripheral tissues was also unaffected (Fig. 1f-i, Extended Data Fig. 2c-e).

## Increased catecholamine, decreased insulin secretion, and enhanced gluconeogenesis during the post-CSDS period

Epinephrine and norepinephrine were elevated during the post-CSDS period but not immediate-CSDS period (Fig. 1j, Extended Data Fig. 2f). As sympathetic nervous system (SNS) is known to increase gluconeogenesis, we evaluated gluconeogenesis by injecting pyruvate, amino acids, and glycerol. Blood glucose levels did not increase in the pyruvate or alanine tolerance tests during the immediate- or post-CSDS period (Extended Data Fig. 3a-d). However, in the glycerol tolerance test, CSDS-exposed mice had higher blood glucose levels than control mice during the post-CSDS but not immediate-CSDS period (Fig. 1k, Extended Data Fig. 3e). We also measured insulin secretion during the post-CSDS period using a hyperglycemic clamp (Fig. 1l-n). The glucose infusion rate in the post-CSDS group was lower than that of the control at the start of the experiment but showed no significant changes during the steady state (Fig. 1m, Extended Data Fig. 3f, g). In post-CSDS mice, C-peptide levels were reduced (Fig. 1n). These results suggest that increased SNS and decreased insulin secretion promote hepatic gluconeogenesis during the post-CSDS period.

## Cerebellar fastigial nucleus is activated in post-CSDS mice

To study what neuronal circuits may enhance gluconeogenesis in CSDS-exposed mice in the post-CSDS period, we measured glucose uptake in various brain regions using 2-[^14^C]-Deoxy-D-Glucose (2-[^14^C] DG). Mice were injected with 2-[^14^C] DG after performing the HE-clamp. Interestingly, increased 2-[^14^C] DG uptake was observed only in the cerebellum (Fig. 1o). Though the cerebellum classically is studied in the context of fine-tuning of motor control and coordination, it has also been found to be significantly involved in fear-related emotional regulation and mood disorders^6–9^. Cerebellar activation has been observed in major depression^6–8^ and PTSD^6,7^ but its relationship to glucose metabolism has never been investigated. To more closely evaluate cerebellar neuronal activity in post-CSDS mice, we recorded the electrical activity of cerebellar cortical neurons using high-density microelectrode arrays (Extended Data Fig. 4a, b). The inter-spike interval (ISI) of post-CSDS mice was similar to that of control mice when brain slices were perfused with aCSF (Extended Data Fig. 4c). However, when brain slices were perfused with a high potassium solution, GABA receptor antagonists, or glutamate receptor antagonists (which increase firing rate), cerebellar cortical neurons of post-CSDS mice did not show the expected decrease in ISI observed in control mice (Extended Data Fig. 4c). These findings suggest that cerebellar cortical neurons exhibit abnormal glutamatergic and GABAergic regulation during the post-CSDS period. We then focused further on neuronal activity within the cerebellar fastigial nucleus (FN), which has been implicated in fear-related emotional regulation^7–9^. Using an in vivo electrophysiological recording in an awake state in control and post-CSDS mice (Fig. 1p), we observed a significantly higher firing rate in FN neurons in the post-CSDS mice compared to those in the control mice (Fig. 1q, r).

## FN neurons regulate anxiety-like behavior and blood glucose levels

To examine the contribution of FN neurons to anxiety-like behavior, we used an excitatory DREADD system (Fig. 2a). Activation of FN neurons in naive mice by DREADD agonist (clozapine, CLZ) induced marked anxiety-like behavior, similar to CSDS-exposed mice in the post-CSDS period (Extended Data Fig. 5a). Activation of FN neurons was also sufficient to promote glucose intolerance and gluconeogenesis, and was also accompanied by an increase in plasma epinephrine (Fig. 2b-d, Extended Data Fig. 5b-e). The activation of FN neurons did not affect insulin sensitivity, insulin secretion, or the levels of corticosterone in the plasma (Extended Data Fig. 5f-h). We next used an inhibitory DREADD system to see if this could rescue CSDS-induced changes (Fig. 2e, Extended Data Fig. 6a). Inhibition of FN neurons during the post-CSDS period acutely improved glucose tolerance and suppressed glycerol-driven gluconeogenesis (Fig. 2f, g). Inhibiting FN neurons also did not affect insulin resistance, insulin secretion or corticosterone levels (Extended Data Fig. 6b-d). Thus, the activation of FN neurons after CSDS is likely responsible for the glucose metabolic abnormalities and anxiety-like behavior during the post-CSDS period.

**Fig. 2.**
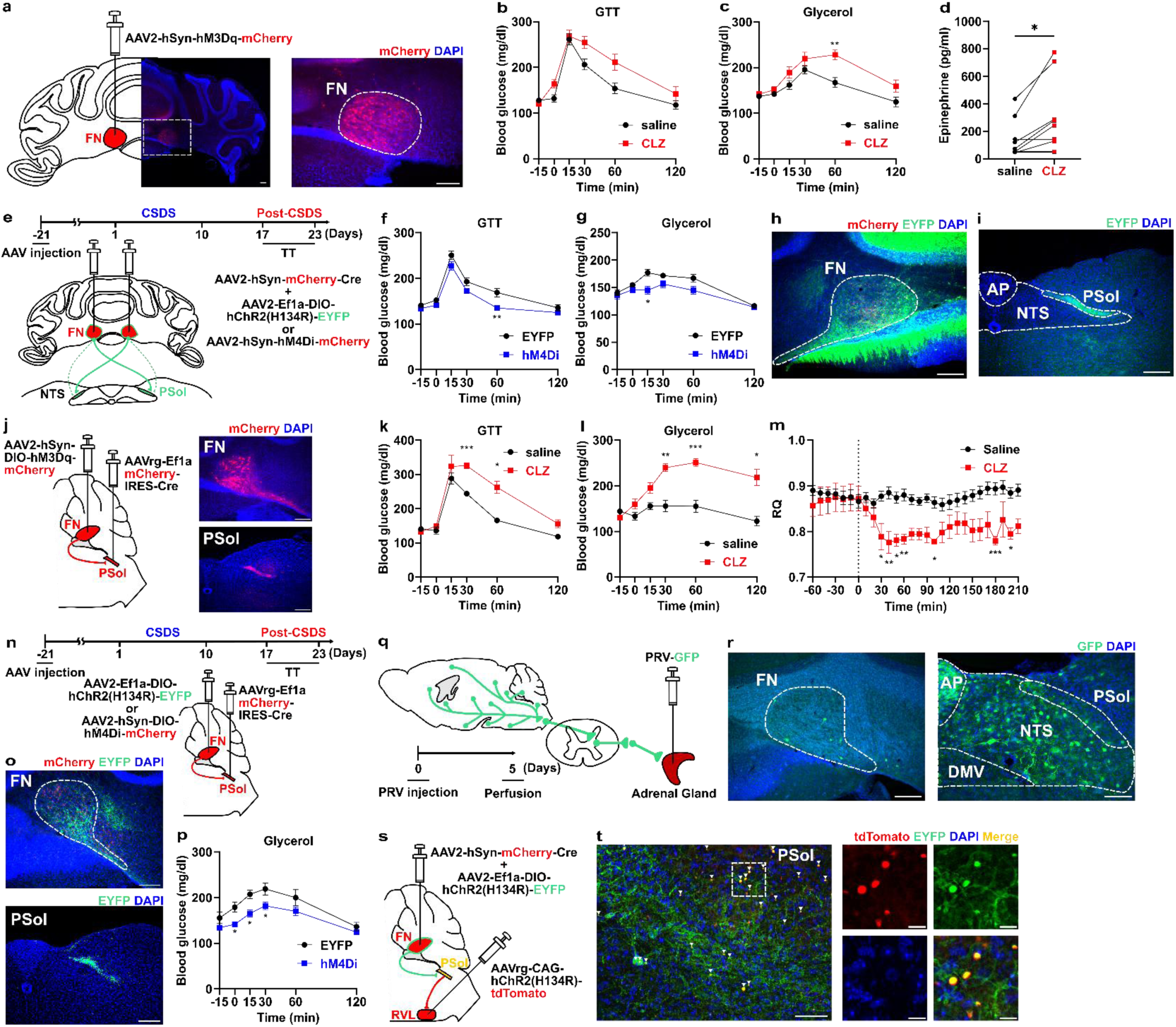
FN neurons projecting to the PSol regulate whole-body metabolism and depression-like behavior. **a**, Schematic of DREADD virus injection and mCherry expression. **b**, Glucose tolerance test (GTT, 0–120 min) after saline or clozapine (CLZ) injection (-15 min) (n = 10). **c**, Glycerol tolerance test (0–120 min) after saline or CLZ injection (-15 min) (n = 9). **d**, Plasma epinephrine levels after saline or CLZ injection (-30 min) (n = 9). **e**, Schematic of virus injections and timeline of experiments. **f**, GTT (0-120 min) after CLZ injection (-15 min) in EYFP (n = 8) or hM4Di mice (n = 6). **g**, Glycerol tolerance test (0–120 min) after CLZ injection (-15 min) in EYFP (n = 10) or hM4Di mice (n = 7). **h**, Expression of mCherry and EYFP in the FN of a control mouse. **i**, EYFP-positive fiber innerved from FN to the parasolitary nucleus (PSol). AP, area postrema; NTS, nucleus tracts solitarius. **j**, Expression of mCherry after virus injections into the FN and PSol. AAVrg, retrograde serotype of AAV. **k**, GTT (0–120 min) after saline or CLZ injection (-15 min) (n = 5). **l**, Glycerol tolerance test (0–120 min) after saline or CLZ injection (-15 min) (n = 5). **m**, Respiratory quotient (RQ) after saline or CLZ injection (0 min) (n = 10). **n**, Schematic of virus injections and timeline of experiments. **o**, mCherry and EYFP expression in the FN (top), EYFP-positive fiber in the PSol (bottom) of a control mouse. **p**, Glycerol tolerance test (0–120 min) after CLZ injection (-15 min) of EYFP (n = 5) or hM4Di mice (n = 9). **q**, Pseudorabies virus (PRV-GFP) was injected into the left adrenal gland. Mice were sacrificed 5 days after PRV injection (n = 5). **r**, Representative PRV-infected regions in the FN and brainstem include PSol. DMV, the dorsal motor nucleus of the vagus. **s,** Schematic of AAV injection into the FN and rostral ventrolateral medulla (RVL). **t**, EYFP-positive fiber from FN and tdTomato-positive cell retrogradely infected from the RVL (left). The boxed areas on the left are magnified on the right images. Scale bars are 200μm in **a**, **h-j**, **o**, **r**, **t** (left), and 10μm in **t** (right). Data are presented as mean ± SEM; \**p* < 0.05, \*\**p* < 0.01, \*\*\**p* < 0.001, two-way ANOVA followed by Sidak multiple comparison test in **b**, **c**, **f**, **g**, **k-m**, **p**; two-tailed paired t-test in **d**.

## FN-PSol neurons regulate whole-body energy metabolism

To further investigate the mechanism by which the FN exerts the observed effects, we first mapped the brain regions to which FN neurons project. Using mice expressing EYFP in FN neurons, we found that FN neurons project to a wide range of nuclei across the brain (Fig. 2e, h, i, Extended Data Fig. 7). In the forebrain, FN neurons projected to the medial septal nucleus, the nucleus of the diagonal band of Broca, the bed nucleus, and the central amygdala, which are involved in pain and emotional processing (Extended Data Fig. 7b, c, g). In the hypothalamus, FN neurons projected to the preoptic area, the dorsomedial hypothalamus, and the lateral hypothalamus (Extended Data Fig. 7d-f). In the thalamus, FN neurons projected to the paraventricular, medial, and parafascicular thalamic nuclei (Extended Data Fig. 7h, i). FN neurons also projected to the ventrolateral periaqueductal gray and the parabrachial nucleus in the pons (Extended Data Fig. 7j-l). Furthermore, FN neurons projected to the vestibular nucleus and various regions in the brainstem, including the olivary nucleus, parasolitary nucleus (PSol), and the rostroventrolateral reticular nucleus (RVL) (Fig. 2i, Extended Data Fig. 7m-o).

To then identify which of these neuronal circuits contributes to metabolic regulation, we used an excitatory DREADD system specifically in FN neurons projecting to the hypothalamus, thalamus, pons, and medulla, regions known to be involved in metabolic control (Fig. 2j, Extended Data Fig. 8). Most of the neural circuits projecting from the FN did not affect glucose tolerance (Extended Data Fig. 8). Activation of FN neurons projecting to the lateral hypothalamus tended to improve glucose tolerance (Extended Data Fig. 8c). However, only activation of FN neurons projecting to the PSol induced glucose intolerance and enhanced gluconeogenesis (Fig. 2j-l, Extended Data Fig. 9a). It also tended to reduce insulin sensitivity, but unaffected insulin secretion (Extended Data Fig. 9b, c). Direct activation of FN-PSol neurons induced a transient reduction in oxygen consumption and energy expenditure after CLZ administration (Extended Data Fig. 9d, e). Furthermore, the activation of FN-PSol neurons decreased the respiratory quotient for over 3 hours, reducing carbohydrate utilization and increasing lipid utilization (Fig. 2m, Extended Data Fig. 9f, g). Conversely, inhibition of FN-Psol neurons using an inhibitory DREDD system suppressed glycerol-driven gluconeogenesis (Fig. 2n-p), but did not affect insulin resistance (Extended Data Fig. 9i). These results suggest that FN-PSol neurons contribute to whole-body energy metabolism by shifting energy utilization from glucose to lipids.

To assess whether FN-Psol neurons also contribute to anxiety-like behavior, we conducted the OF test in mice with DREDD-activated FN-Psol neurons. Administration of DREDD agonist reduced the time spent in the center and total distance traveled during the OF test, while increasing immobility time (Extended Data Fig. 9h). Thus, activation of FN-PSol neurons is sufficient to induce anxiety-like behavior, similar to the post-CSDS period.

## FN neurons functionally innervate the adrenal glands

Our findings suggest that FN neurons regulate whole-body energy metabolism and endocrine secretion. To assess whether this may be mediated through direct innervation of the adrenal gland, we infected the adrenal glands with pseudorabies virus expressing GFP (PRV-GFP) as a retrograde tracer (Fig. 2q, Extended Data Fig. 10a). GFP expression was observed in widespread brain regions including the FN and PSol (Fig. 2r, Extended Data Fig. 10b-y). Among cerebellar nuclei, PRV-labeled neurons were only found in the FN (Extended Data Fig. 10u). Epinephrine secretion from the adrenal medulla is known to be innervated by the RVL^18,19^. We investigated whether FN neurons that project to the PSol have a connection with the RVL (Fig. 2s). EYFP expressed in FN neurons and tdTomato expressed retrogradely from the RVL were co-localized in PSol neurons (Fig. 2s, t), suggesting that FN neurons functionally innervate the adrenal medulla via the PSol and RVL.

## Single-cell analysis of deep cerebellar nuclei (DCN) cells following CSDS exposure

To investigate changes in the DCN, including FN, cell population after CSDS, we performed single-cell (sc) RNA sequencing. DCN cells were collected from control, immediate-CSDS, and post-CSDS mice, and clustering was performed after initial data processing (Fig. 3a, b). The population of myeloid-derived suppressor cells (MDSCs) was increased in the post-CSDS (Fig. 3a). We next used CellChat to infer intercellular communications underlying the effect of CSDS exposure on neurons. Several cell types may affect neurons via Negr1-Negr1 signaling (Fig. 3c), but we found that the communication between astrocytes and neurons via Negr1 signaling was markedly decreased by CSDS exposure (Extended Data Fig. 11a).

**Fig. 3.**
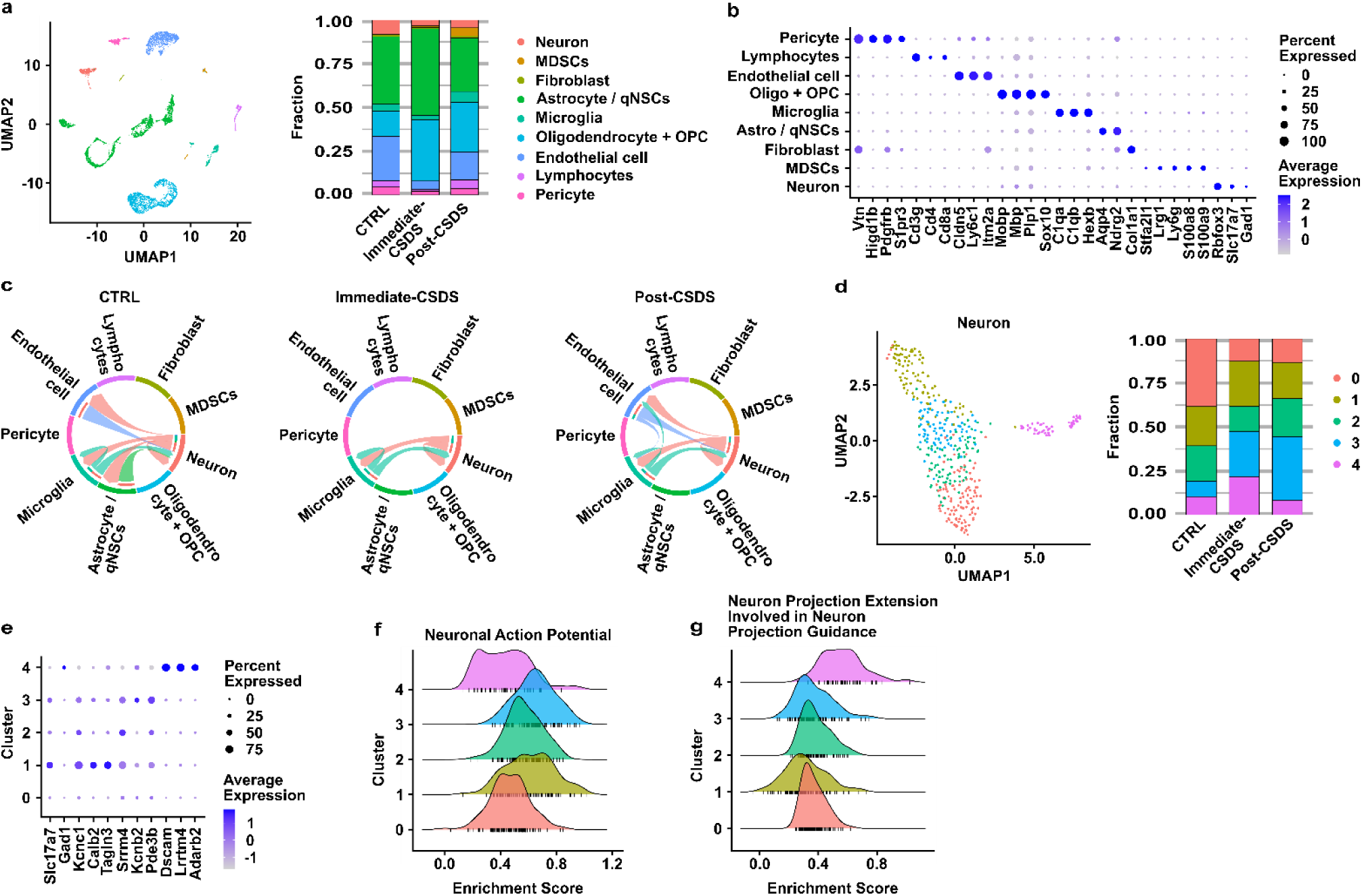
Single cell analysis of DCN following CSDS exposure. **a**, Uniform Manifold Approximation and Projection (UMAP) plot of 7,747 cells from control (2,832 cells), immediate-CSDS (1,600 cells), and post-CSDS (3,158 cells) mice (left). Different colors represent different cell populations. The proportion of cells in each cluster is shown for each group (right). MDSCs, myeloid-derived suppressor cells; qNSCs, quiescent neural stem cells; OPC, oligodendrocyte precursor cells. **b**, Expression of the indicated marker genes across different cell types. **c**, Circus plot illustrating cellular crosstalk via Negr1-Negr1. **d**, UMAP plot of 426 neurons (left). The proportion of neurons in each of the five clusters is shown for each group (right). **e**, Expression of the indicated marker genes across different neuronal cell types. **f**, **g**, Gene set enrichment analysis (GSEA) showing GO terms with increased enrichment scores in cluster 3 (**f**) and cluster 4 (**g**).

We next clustered neuronal cells and generated 5 subclusters (C0-C4) (Fig. 3d). Gene enrichment analyses suggest that C0 might be involved in immune response, C1 in the regulation of neuronal synaptic plasticity, C2 in the regulation of feeding behavior (Extended Data Fig. 11b-d). C1-3 expressed Kcnc1 and/or Calb2, which have been reported to be expressed in FN neurons^10^. Notably, C3 cells increased during the post-CSDS period and were linked to neuronal excitability (Fig. 3d-f), suggesting C3 as a neuronal population involved in anxiety-like behavior and whole-body metabolic regulation during the post-CSDS period. C4 cells also increased immediately after CSDS exposure (Fig. 3d). These cells express Dscam and Lrrtm4 and were associated with neural circuit formation (Fig. 3e, g). Thus, CSDS exposure may change axon guidance and synapse formation in C4 neurons, potentially contributing to anxiety-like behavior and escape behavior.

## Depression is associated with cerebellar change and glucose intolerance in human patient data

To assess if any evidence for functional connectivity between the cerebellum, mood disorders, and diabetes exists in humans, we first assessed the relationship between cerebellar volume, symptoms of depression, and diabetes in a large-scale cohort of 1325 patients in Arao, Kumamoto, Japan (the Arao cohort). The volume of the cerebellar white matter, including the FN, showed a weak correlation with the Geriatric Depression Scale (GDS) (Fig. 4a, b), but a significant correlation with HbA1C which is stronger in participants with depression (Fig. 4c, d). To better characterize these relationships, we made a generalized linear mixed model relating cerebellar white matter volume, HbA1C, and GDS (Fig. 4e). HbA1C had a significant main effect on cerebellar white matter volume (F (1316) = 6.64, p = 0.01), whereas GDS type alone was not significant (F (1316) = 0.58, p = 0.56). However, a significant interaction between GDS type and HbA1C was observed, suggesting that the effect of GDS type on cerebellar white matter volume depends on the level of HbA1C.

**Fig. 4.**
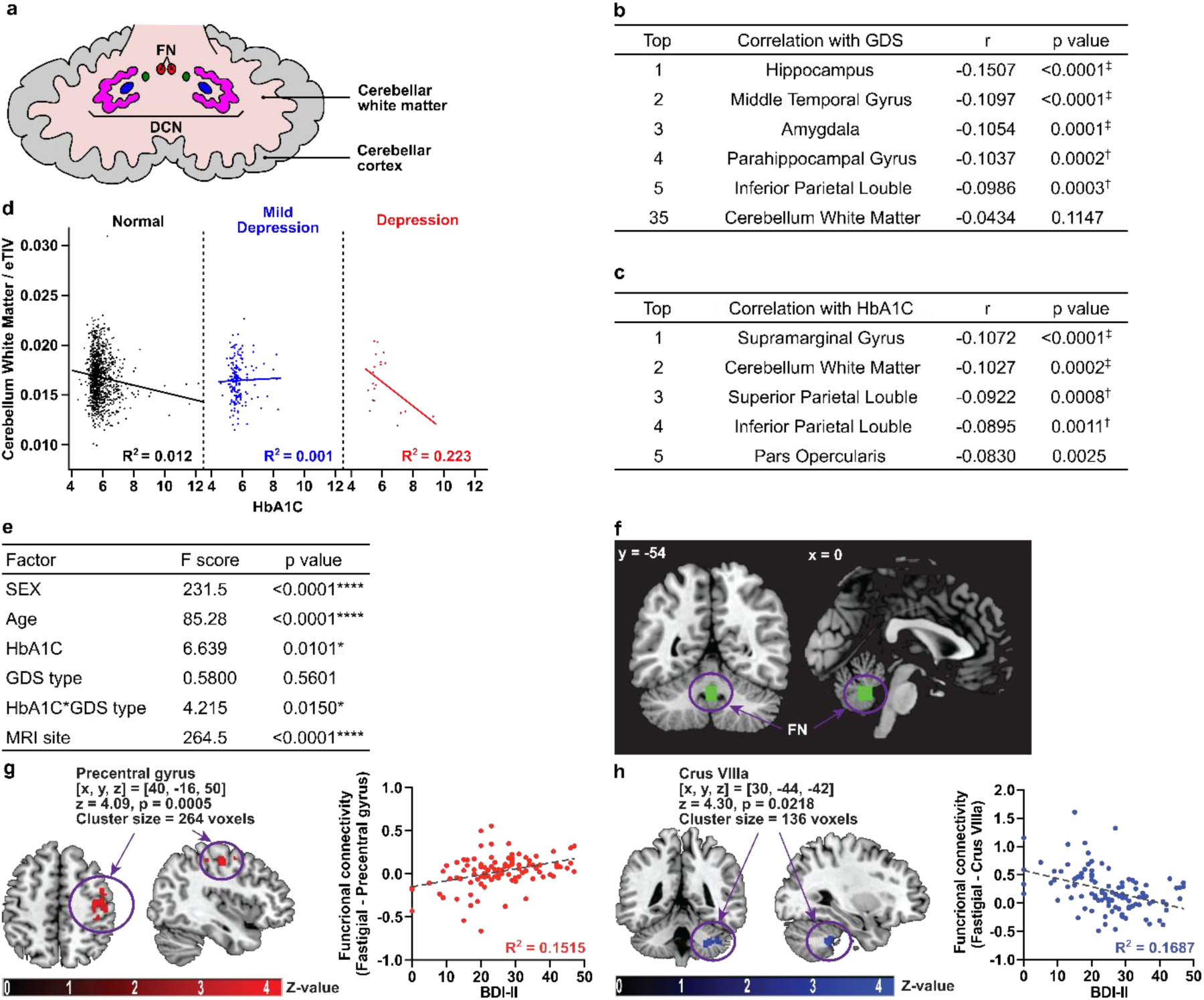
Cerebellar abnormalities correlate with depressive symptoms and hyperglycemia. **a**, Schematics of cerebellar anatomy. The cerebellum consists of the cerebellar cortex and cerebellar white matter. The cerebellar white matter contains the deep cerebellar nucleus (DCN), including the fastigial nucleus (FN). **b-e**, Studies for the Arao cohort (n = 1325). **b**, Correlation between the Geriatric Depression Scale (GDS) and brain region volumes. The top five regions and cerebellar white matter are shown. **c**, Correlation between HbA1C and brain region volumes. **d**, Based on GDS scores, subjects were grouped into normal (GDS 0–4, n = 1140), mild depression (GDS 5–9, n = 165), and depression (GDS 10–15, n = 20). In participants with depression, cerebellar white matter volume showed a stronger correlation with HbA1C. **e**, A generalized linear mixed model (GLMM) was used to analyze the effects of each factor on cerebellar white matter volume as the dependent variable. **f-h**, Functional connectivity of the FN and other brain regions using fMRI data from individuals with major depressive disorder. **f**, Anatomical mask of FN used in this analysis is shown as a green square. **g**, Axial and sagittal sections of greater functional connectivity between the FN and the pre-central gyrus in the patients with depression (left panel). Higher functional connectivity is correlated with the higher Beck Depression Inventory (BDI-II) scores (*p* = 0.0005) (right panel). **h**, Axial and sagittal sections of greater functional connectivity between the FN and the cerebellar tonsil (left panel). Functional connectivity of this pathway is negatively correlated with BDI-II scores (*p* = 0.0218) (right panel). †*p* < 0.0011, ‡*p* < 0.0002, the *p*-values were calculated using Pearson correlation, and Bonferroni correction was applied to account for 44 multiple comparisons in **b**, **c**: \**p* < 0.05, \*\*\*\**p* < 0.0001, the F-test in **e**.

Next, we evaluated how depression affects functional connectivity of cerebellum. Using fMRI data from 104 patients diagnosed with major depressive disorder (MDD), we correlated measures of functional connectivity of the FN with Beck Depression Inventory (BDI-II) scores^11^. Patients with higher BDI-II scores had stronger functional connectivity between the FN and the pre-central gyrus (Fig. 4f) and weaker functional connectivity between the FN and the cerebellar tonsil (Fig. 4g). These findings suggest that depression is associated with altered FN connectivity.

## Discussion

In this study, we characterize in detail the impact of CSDS on systemic glucose metabolism, and describe a novel role of the cerebellar FN in mediating both the observed metabolic and behavioral changes in the post-CSDS period. We find impaired glucose tolerance in the post-CSDS period is associated with increased blood catecholamines and decreased insulin secretion, and is likely independent of HPA activity given low corticosterone levels in this period. We instead note increased activation of the cerebellar FN after CSDS, and identify a FN-Psol pathway activated during the post-CSDS period that is both necessary and sufficient to induce glucose intolerance and anxiety-like behaviors. Moreover, we observe that activation of FN-Psol neurons shifts the body’s primary energy source from glucose to lipids. By single-cell RNA sequencing, we further characterize the impact of CSDS on neuronal connectivity and immune cell interactions in the DCN. We also show that activation of the FN after CSDS results in epinephrine secretion via direct projections to the adrenal glands. Finally, analyses of human patient demographic and fMRI data show correlations between cerebellar white matter volume with HbA1c and depression scores, as well as alterations in FN connectivity in patients with more severe depression.

To our knowledge, this is the first report highlighting the critical role of the cerebellum in regulating systemic metabolism. It also uniquely sheds new light into neuroendocrine pathways involved in chronic or delayed responses to psychological stressors, which up to this point were unknown. While psychological stress is well-known to acutely increase cortisol and catecholamine secretion^12^ via pathways involving the PFC, PVH, VMH, LC, RPa, and intermediolateral nucleus^13^, conditions such as depression^14^, bipolar disorder^15^, and PTSD^16,17^ cause long-term changes in plasma catecholamines and cortisol levels^18–20^ which are better modeled by our delayed post-CSDS model system.

Though the cerebellum is known to regulate motor control and learning^8^, it also plays key roles in prediction^8^, fear-related emotions^9^, psychiatric disorders^6,21^, cardiovascular system^22,23^, feeding behavior^24^, and insulin secretion^25^. The FN has also been described to modulate blood pressure via the autonomic nervous system^26^, thus it’s observed neuroendocrine role in the post-CSDS period was not completely unexpected. Consistent with previous studies^27,28^, PRV-labeled neurons from the adrenal gland were identified in neural nuclei involved in sympathetic nervous system control, stress and pain response, and metabolic regulation^29^, and FN. PRV-labeled neurons were also observed in the motor cortex, primary somatosensory cortex, parietal association cortex, and other regions of the neocortex responsible for processing sensory information such as vision, hearing, olfaction, and spatial awareness (Extended Data Fig. 10h, i, k). In our fMRI study, more robust functional connectivity between the FN and the pre-central gyrus, including in the primary motor cortex in mice, was associated with higher BDI-II scores. This connection may provide new insights into stress responses centered on the cortex-adrenal gland pathway.

Our observation that FN projections covered a wide range of brain regions, including the cerebrum, hypothalamus, thalamus, PAG, pons, and medulla, is consistent with that of prior studies^8,10,30^. FN-PSol neurons in particular have been suggested to be involved in integration of sensorimotor and autonomic information^31^. PSol projections to the NTS and RVL have also been previously reported^32^. We confirm that the PSol connects the pathway between FN and RVL, where sympathetic preganglionic neurons to the adrenal gland exist^33^. The specific peripheral organs that are innervated by FN-PSol neurons remain unclear.

Our scRNA sequencing analysis further improves our understanding of how the DCN is remodeled after CSDS and the role of specific DCN neurons. In particular, we identify a neuronal population involved in axon guidance that increased immediately after CSDS (C4) and another population that exhibited excitatory properties during the post-CSDS period (C3), which may be responsible for anxiety-like behavior and whole-body metabolic regulation in the FN. Additionally, the observed decrease in C0, which expresses genes associated with immunosuppression, and increase in MDSCs during the post-CSDS period (Extended Data Fig. 11b, Fig. 3d) suggest that CSDS may alter immune regulation in the DCN as well. The characterization of these neuronal clusters distinct from those associated with motor function and neuronal plasticity holds promise for advancing future cerebellar research.

Finally, our observed correlations between cerebellar white matter volume and functional connectivity with HbA1c and depression scores corroborate prior studies noting increased cerebellar activity in patients with depression and PTSD^6–8^. It is also supported by the observation that patients with depression^2^ and PTSD^3,4^ have a higher risk of developing diabetes. Our study links the importance of the cerebellum in psychiatric disorders to the development of diabetes. Advancing research on cerebellar regulation of metabolism is expected to deepen our understanding of systemic metabolic control and contribute to developing effective therapies for psychiatric disorders and diabetes.

---

**Extended Data Fig. 1.**
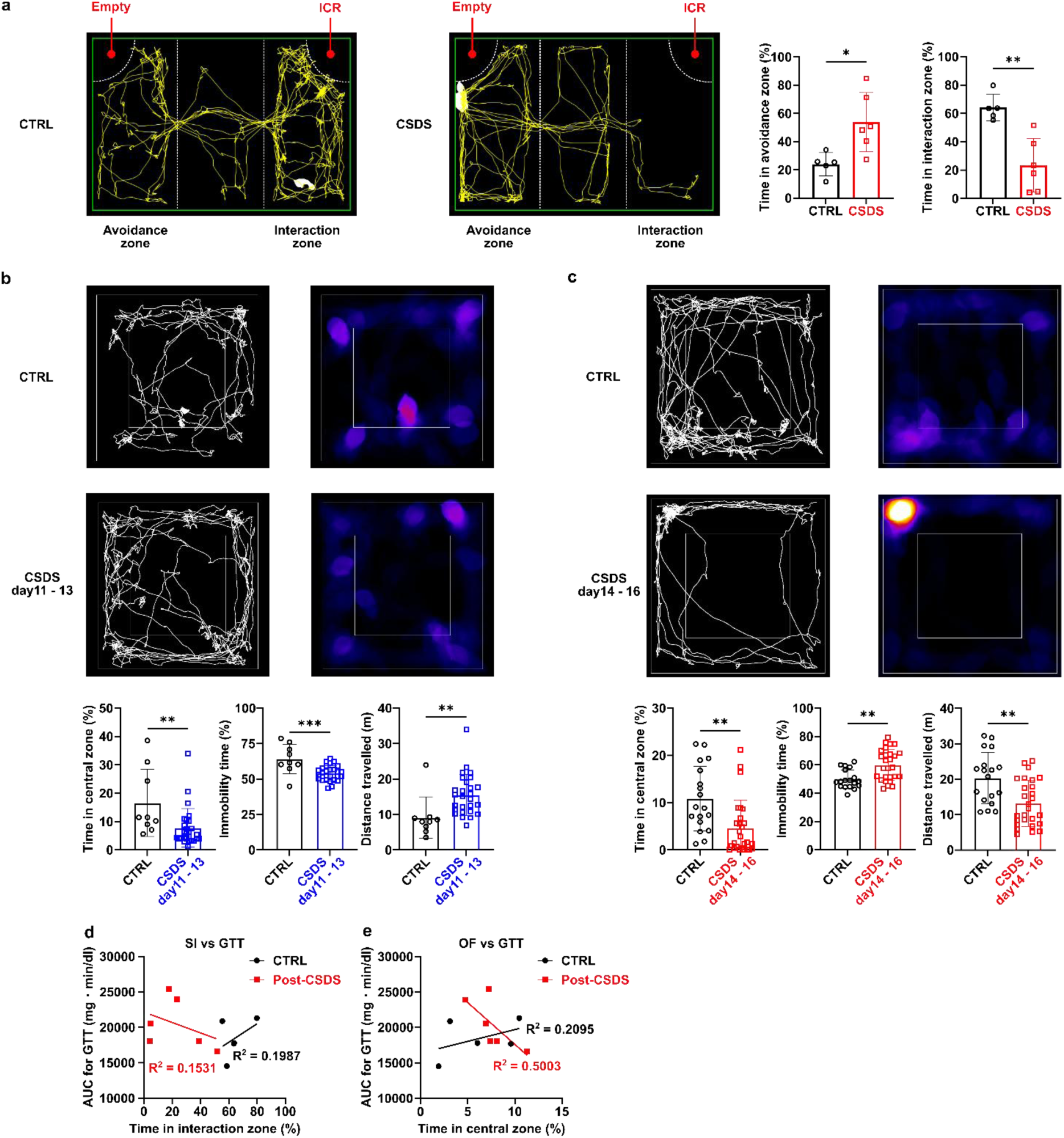
CSDS induces depression-like behavior. **a**, Social interaction test (SI) was performed using a three-chambered system to evaluate social avoidance. The interaction zone contained a cage with an ICR mouse, while the avoidance zone contained an empty cage. On day 13, CSDS-exposed mice spent more time in the avoidance zone and less time in the interaction zone (CTRL, n=5; CSDS, n=6). **b**, **c**, Open field test (OF) in CSDS day 11-13 (**b**) and day14-16 (**c**). On day 11-13, the time spent in the center was reduced, while the total distance traveled increased (**b**, CTRL, n=9; CSDS, n=27). On day 14-16, the total distance traveled decreased, and immobility time increased (**c**, CTRL, n=18; CSDS, n=27), suggesting that depression-like behavior strengthens on day 14-16. **d**, **e**, The correlation between the results of the SI (**d**) or OF (**e**) and glucose tolerance test (GTT) (CTRL, n=5; CSDS, n=6). Data are presented as mean ± SEM; \**p* < 0.05, \*\**p* < 0.01, \*\*\**p* < 0.001, two-tailed t-test in **a-c**.

**Extended Data Fig. 2.**
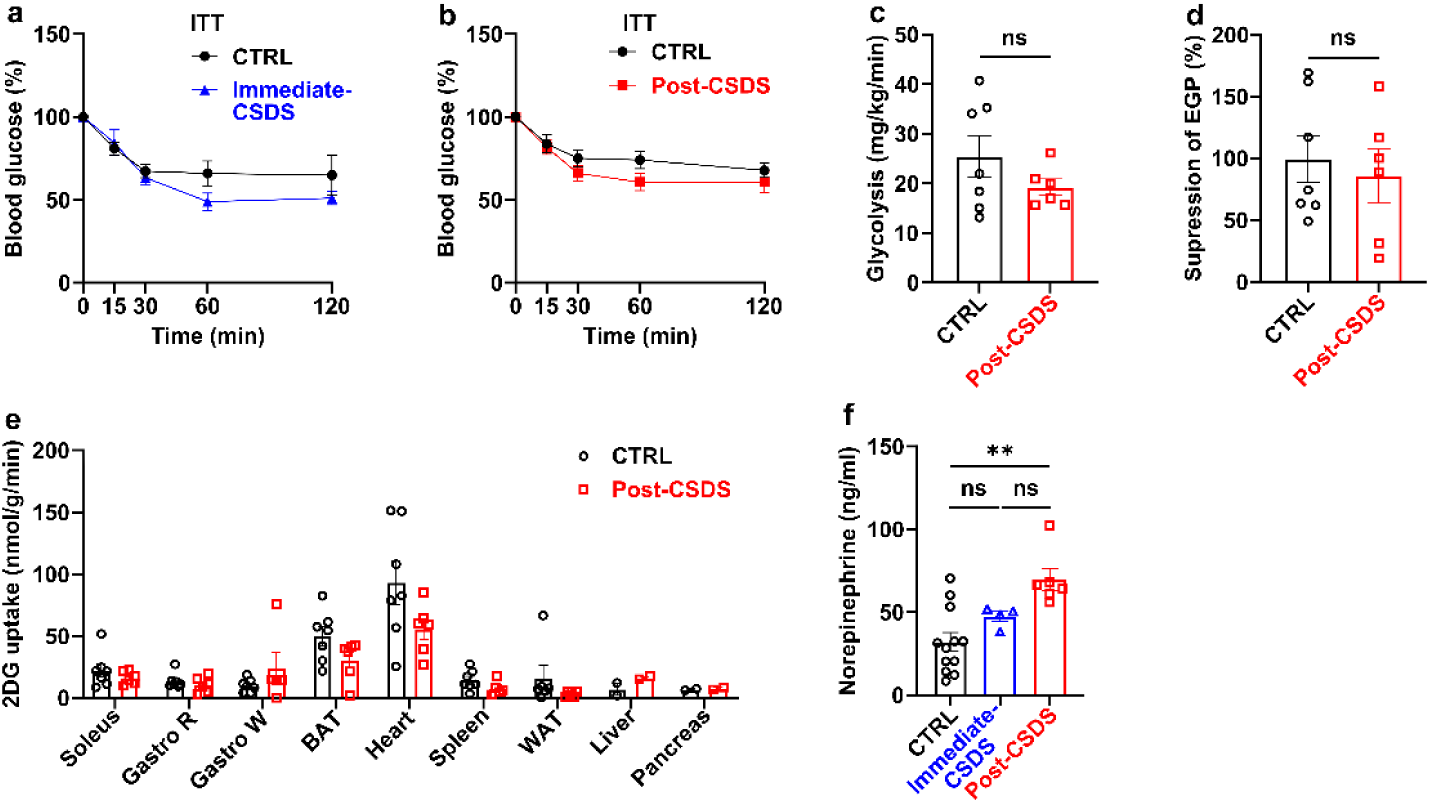
CSDS exposure does not affect insulin sensitivity. **a**, Insulin tolerance test (ITT) of control (CTRL, n = 6) or immediate-CSDS mice (n = 6). **b**, ITT of control (n = 10) or post-CSDS mice (n = 10). **c-e**, Hyperinsulinemic-euglycemic clamp (HE Clamp) studies in control (n = 7) and post-CSDS (n = 6) mice. **c**, The rates of whole-body glycolysis in control and post-CSDS. **d**, Insulin-induced suppression of endogenous glucose production (EGP), which represents hepatic insulin sensitivity in control and post-CSDS. **e**, 2-[^14^C]-Deoxy-D-Glucose (2DG) uptake in soleus, red-portion of the gastrocnemius muscle (Gastro R), brown adipose tissue (BAT), and heart (CTRL, n = 7, Post-CSDS, n = 6); white-portion of the gastrocnemius muscle (Gastro W) and spleen (CTRL, n = 7, Post-CSDS, n = 5); white adipose tissue (WAT) (CTRL, n = 6, Post-CSDS, n = 6); liver and pancreas (CTRL, n = 2, Post-CSDS, n = 2). **f**, Plasma norepinephrine concentration of control, immediate-CSDS, and post-CSDS mice (n = 12, 4, 6). Data are presented as mean ± SEM; \**p* < 0.05, \*\**p* < 0.01, two-way ANOVA followed by Sidak multiple comparison test in **a** and **b**; two-tailed t-test in **c-e**; one-way ANOVA followed by Tukey’s multiple comparison test in **f**.

**Extended Data Fig. 3.**
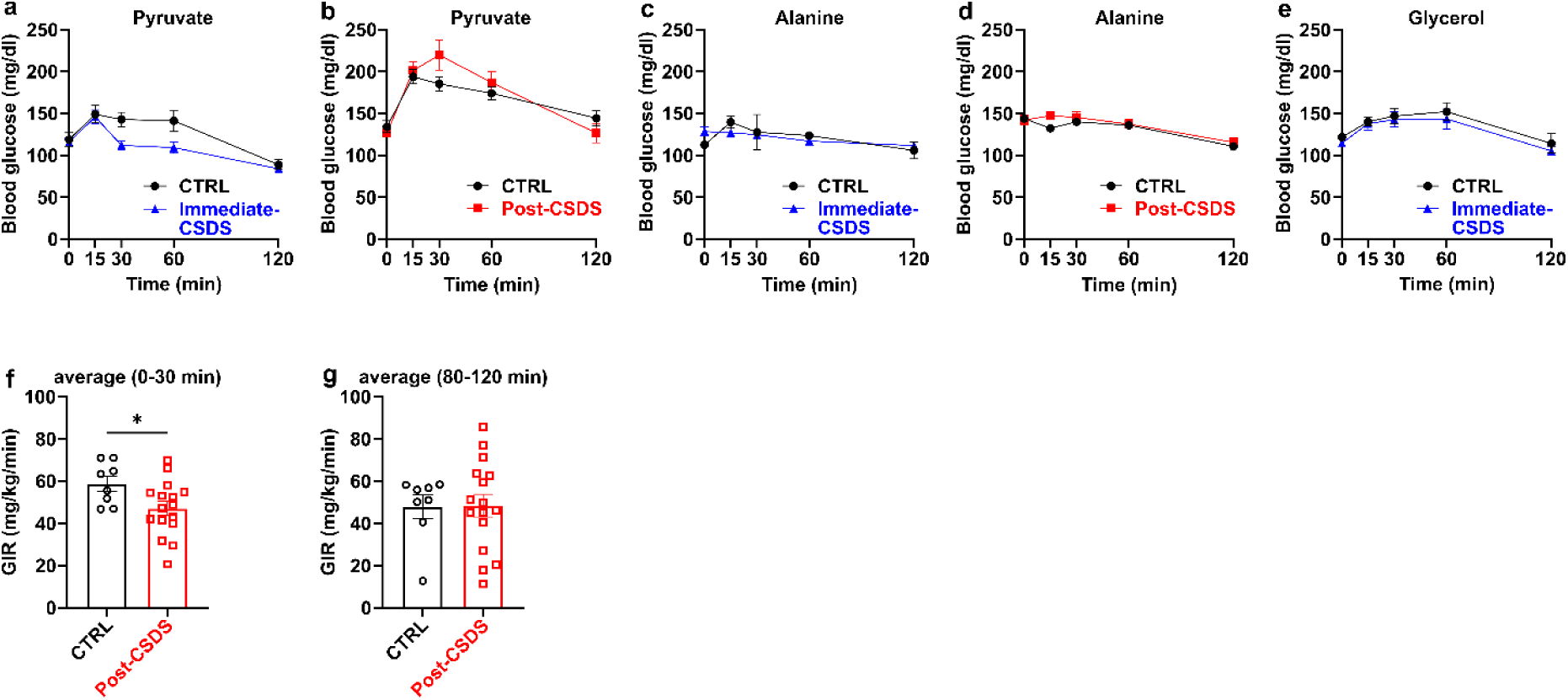
Only glycerol-driven gluconeogenesis is enhanced during the post-CSDS period. **a**, Pyruvate tolerance test of control (CTRL, n = 4) or immediate-CSDS mice (n = 6). **b**, Pyruvate tolerance test of control (n = 8) or post-CSDS mice (n = 6). **c**, Alanine tolerance test of control (n = 3) or immediate-CSDS mice (n = 3). **d**, Alanine tolerance test of control (n = 8) or post-CSDS mice (n = 9). **e**, Glycerol tolerance test of control (n = 5) or immediate-CSDS mice (n = 9). **f**, **g**, Average of glucose infusion rate (GIR) between 0 - 30 min (**f**) and 80 - 120 min (**g**) in hyperglycemic clamp, related to Fig. 1m. GIR is low in post-CSDS mice in 0-30 min. Data are presented as mean ± SEM; \**p* > 0.05, two-way ANOVA followed by Sidak multiple comparison test in **a-e**; two-tailed t-test in **f** and **g**.

**Extended Data Fig. 4.**
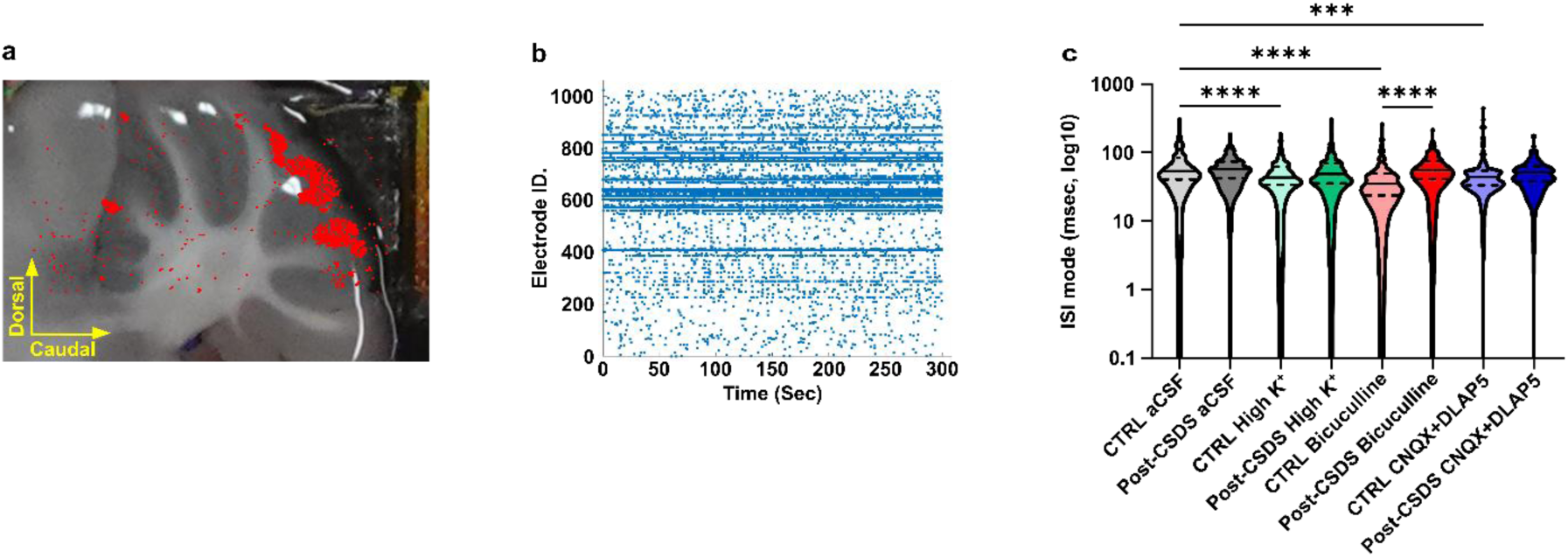
Electrophysiological analysis of cerebellar cortical neurons using a High-Density Microelectrode Array (HD-MEA). **a**, A representative cerebellar recording site. Red spot shows the place where electrical signals were recorded. **b**, Representative examples of recorded neuronal activity in each cell. **c**, Changes in the inter-spike interval (ISI) in cerebellar slices from control and post-CSDS mice after perfusion with high potassium (High K^+^), GABA receptor antagonists (Bicuculline), and glutamate receptor antagonists (CNQX + DL-AP5). \*\*\**p* < 0.001, \*\*\*\**p* < 0.0001, one-way ANOVA followed by Sidak multiple comparison test in **c**.

**Extended Data Fig. 5.**
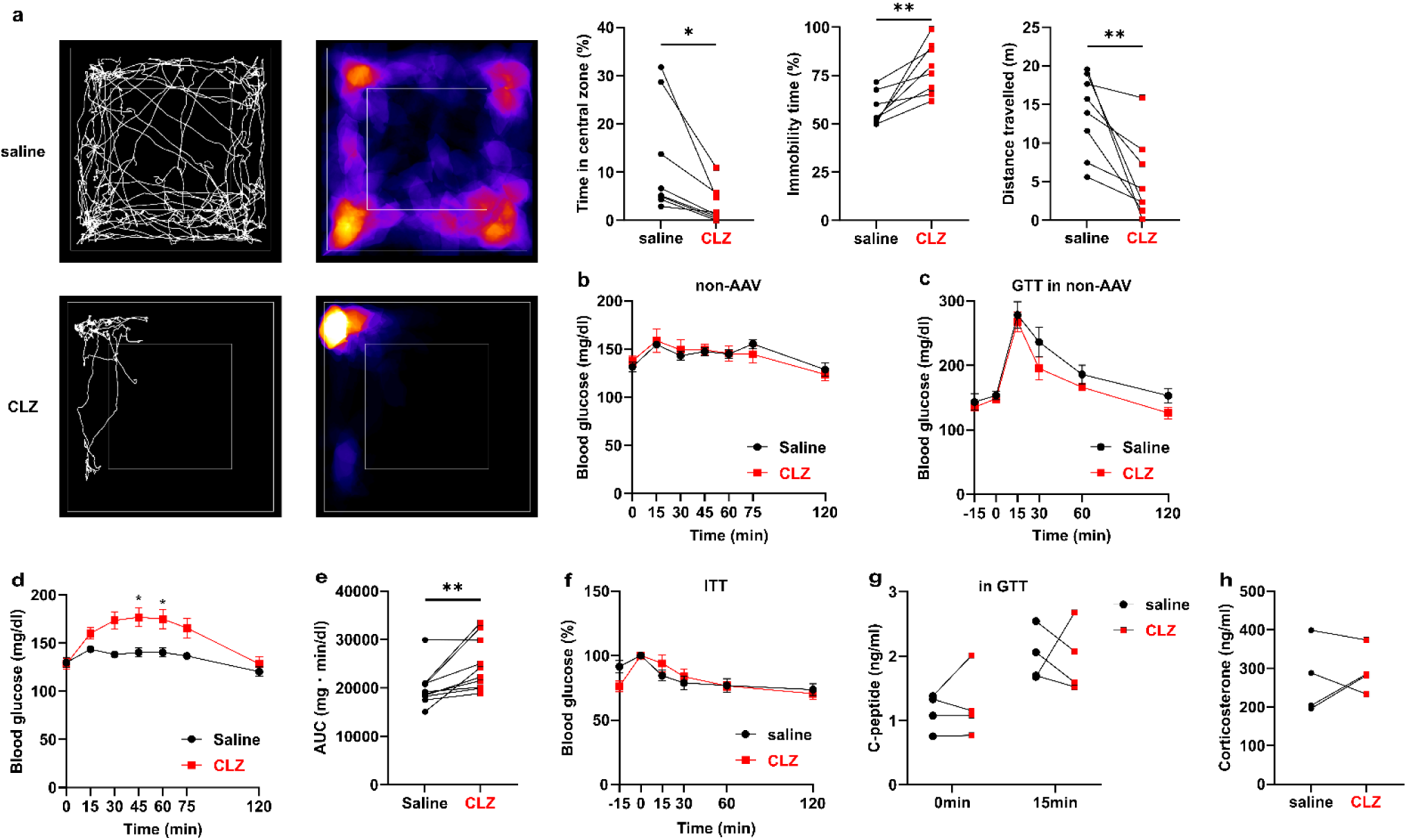
Activation of FN neurons induces anxiety-like behavior and enhances gluconeogenesis. **a**, Open Field test after saline or clozapine (CLZ) injection (-15 min) into the mice expressing excitatory DREADD in FN (n = 8). Mice were injected with AAV2-hSyn-hM3Dq-mCherry in the fastigial nucleus (FN). **b**, Blood glucose levels after the injection of CLZ in control mice (no-AAV injected, n = 5). **c**, GTT (0–120 min) after saline or CLZ injection (-15 min) into the control mice (n = 5). Administration of CLZ to no-AAV-injected mice did not affect blood glucose levels. **d**, Blood glucose levels after saline or CLZ injection (0 min) into the mice injected with AAV2-hSyn-hM3Dq-mCherry in FN (n = 10). **e**, Area under the curve (AUC) of GTT in Fig. 2b. **f-h**, ITT (**f**), plasma C-peptide concentration (**g**), and plasma corticosterone concentration (**h**) after saline or CLZ injection (-15 min) into the mice injected with AAV2-hSyn-hM3Dq-mCherry in FN (**f**, n = 9; **g**, n = 4; **h**, n = 4). Data are presented as mean ± SEM; \**p* < 0.05, \*\**p* < 0.01, paired t-test in **a**, **e**, **g**, and **h**; two-way ANOVA followed by Sidak multiple comparison test in **b-d** and **f**.

**Extended Data Fig. 6.**
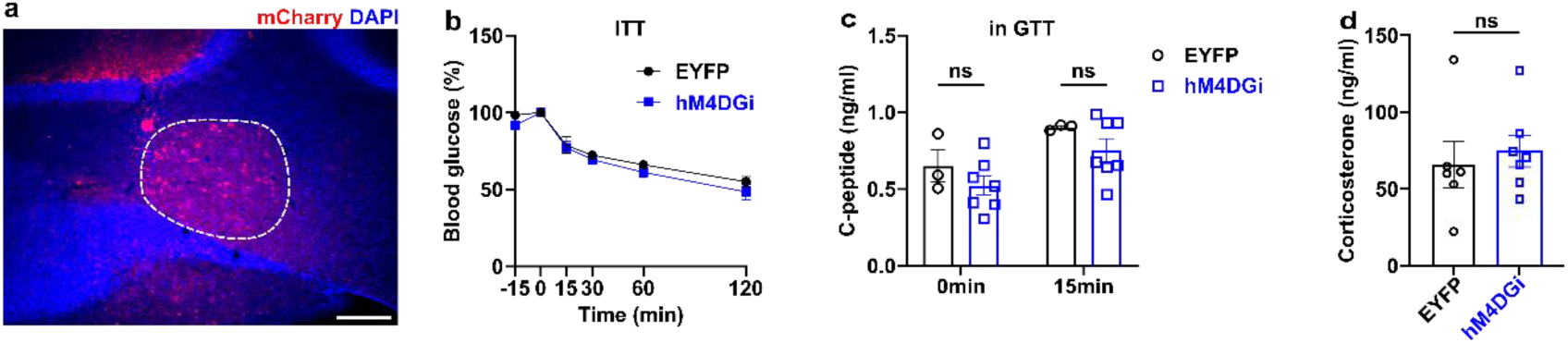
Inhibition of FN neurons during the post-CSDS period does not affect insulin sensitivity, insulin secretion, or corticosterone secretion. **a**, mCherry derived from DREADD virus injected into the FN. The scale bar, 200μm. **b**, ITT (0–120 min) after CLZ injection (-15 min) of EYFP (n = 3) or hM4Di mice (n = 7) in post-CSDS period. **c**, Plasma C-peptide concentration after CLZ injection (-15 min) of EYFP (n = 3) or hM4Di mice (n = 7) in GTT during post-CSDS period. **d**, Plasma corticosterone concentration after CLZ injection (-30 min) of EYFP (n = 6) or hM4Di mice (n = 7). Data are presented as mean ± SEM; \**p* > 0.05, two-way ANOVA followed by Sidak multiple comparison test in **b**; one-way ANOVA followed by Sidak multiple comparison test in **c**; two-tailed t-test in **d**.

**Extended Data Fig. 7.**
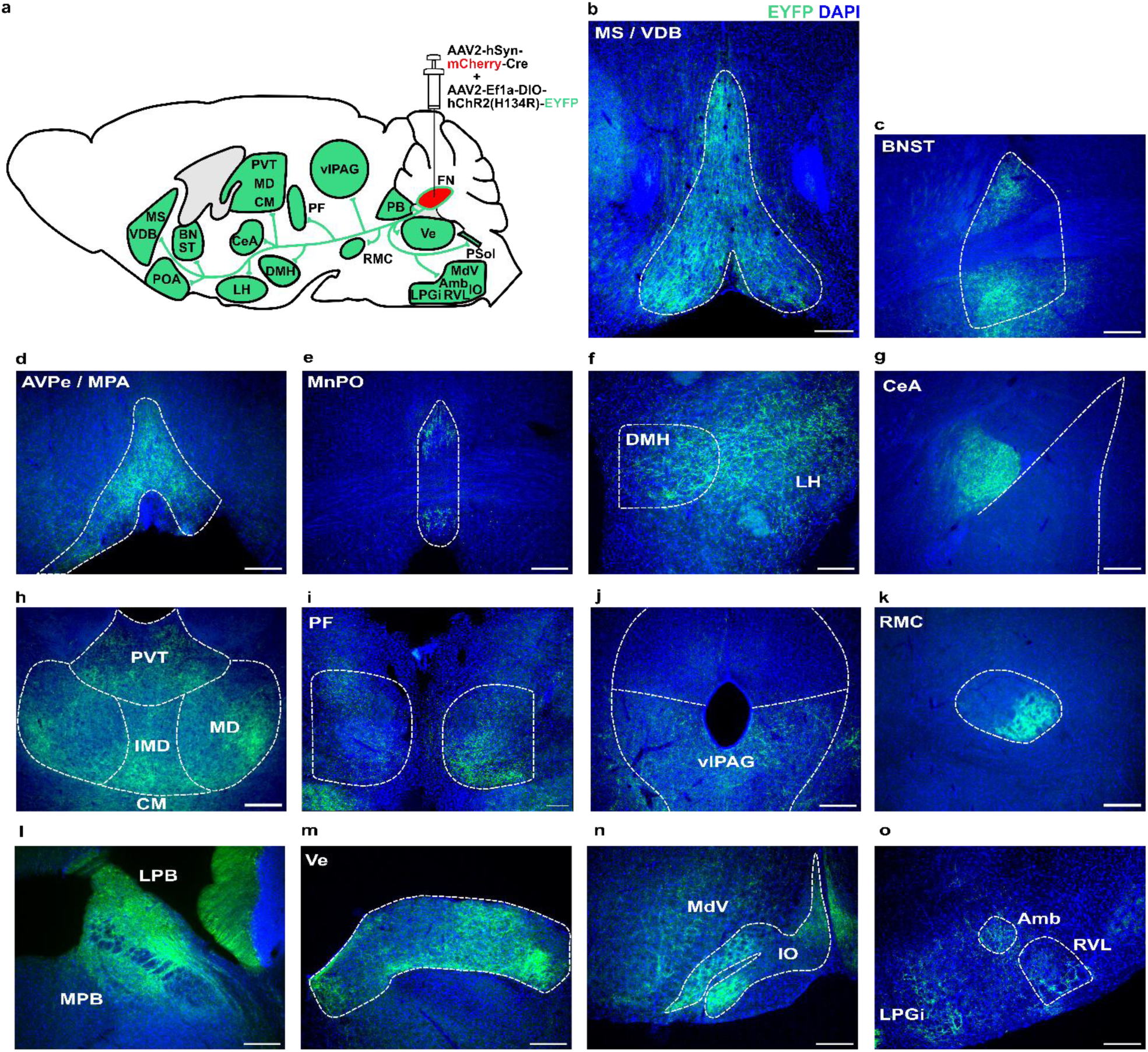
FN neurons project to various regions of the brain. **a**, Schematic of AAV injection into the FN and summary of projection site. **b-o**, EYFP-positive fiber originated from FN. **b**, MS, medial septal nucleus; VDB, the nucleus of the vertical limb of the diagonal band. **c**, BNST, bed nucleus of the stria terminalis. **d**, AVPe, anteroventral periventricular nucleus; MPA, medial preoptic area. **e**, MnPO, median preoptic nucleus. **f**, DMH, dorsomedial hypothalamic nucleus; LH, lateral hypothalamus. **g**, CeA, central amygdaloid nucleus. **h**, PVT, paraventricular thalamic nucleus; MD, mediodorsal thalamic nucleus; IMD, intermediodorsal thalamic nucleus; CM, central medial thalamic nucleus. **i**, PF, parafascicular thalamic nucleus. **j**, vlPAG, ventrolateral periaqueductal gray. **k**, RMC, red nucleus, magnocellular part. **l**, LPB, lateral parabrachial nucleus; MPB, medial parabrachial nucleus. **m**, Ve, vestibular nucleus. **n**, MdV, medullary reticular nucleus ventral part; IO, inferior olive. **o,** LPGi, lateral paragigantocellular nucleus; Amb, ambiguous nucleus; RVL, rostral ventrolateral medulla. Scale bars are 200μm.

**Extended Data Fig. 8.**
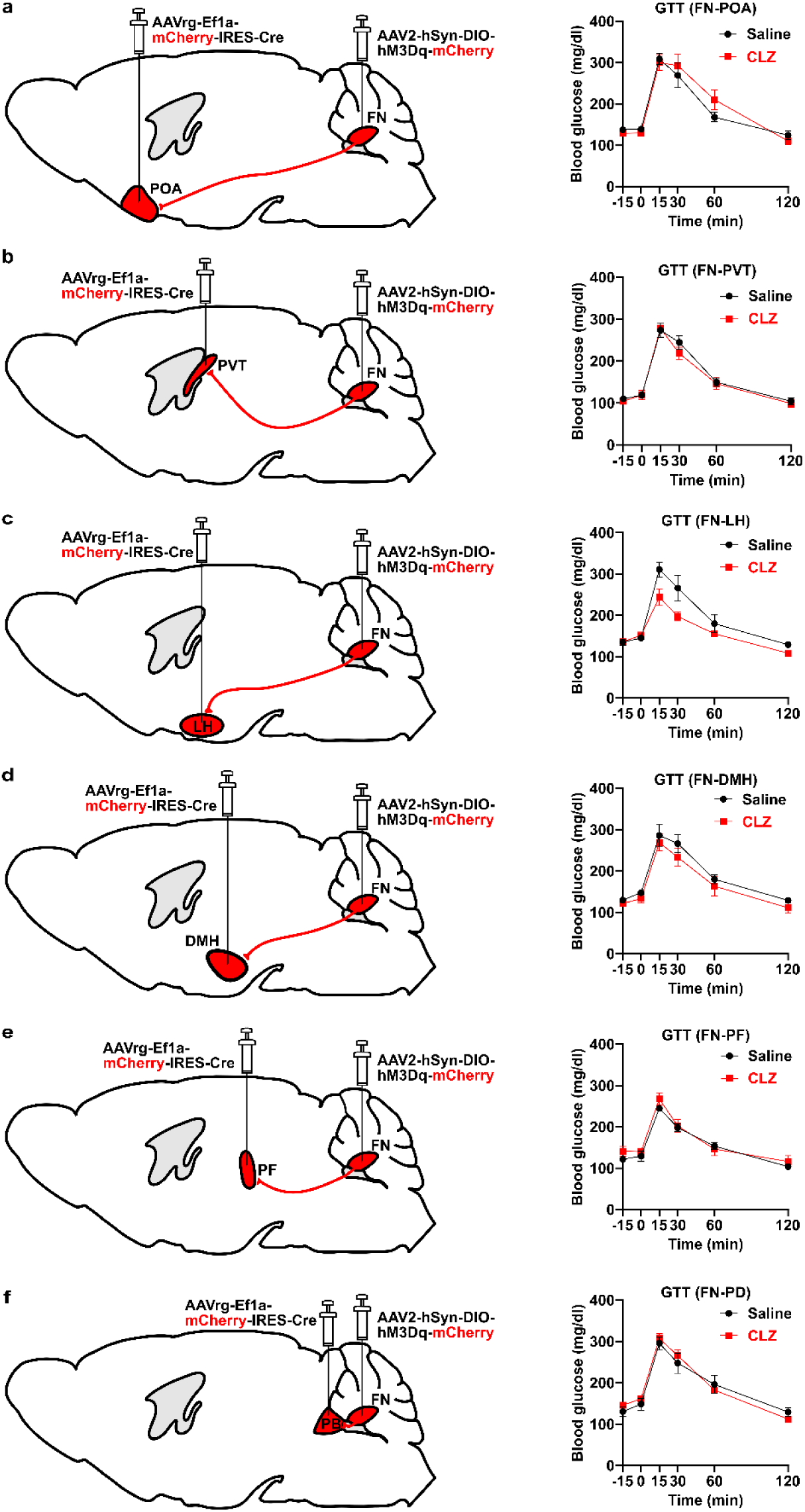
GTT after activating neurons projecting from the FN to each brain region. **a-f**, Schematic of AAV injections and blood glucose levels during GTT **a**, Activation of FN – POA (preoptic area) neurons (n = 5). **b**, Activation of FN – PVT neurons (n = 4). **c**, Activation of FN – LH neurons (n = 4). **d**, Activation of FN – DMH neurons (n = 4). **e**, Activation of FN – PF (parafascicular thalamic nuclus) neurons (n = 4). **f**, Activation of FN – PB (parabrachial nucleus) neurons (n = 3). Data are presented as mean ± SEM; *p* > 0.05, two-way ANOVA followed by Sidak multiple comparison test in **a-f**.

**Extended Data Fig. 9.**
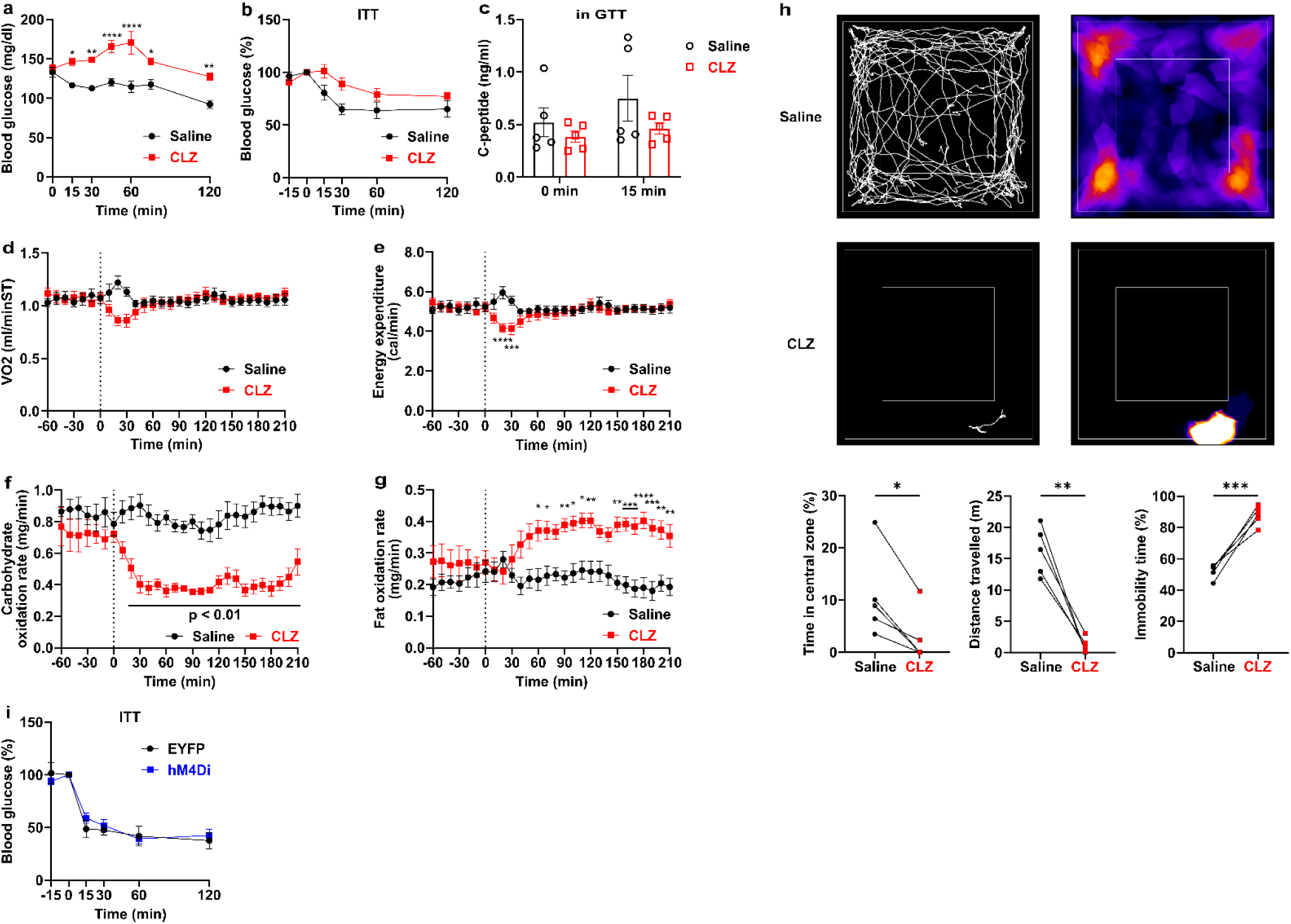
FN-PSol neurons regulate whole-body energy metabolism and anxiety-like behavior. **a-h**, An excitatory DREADD receptor was specifically expressed in neurons projecting from the FN to PSol. **a**, Blood glucose levels after saline or CLZ injection (0 min) into the mice (n = 5). Activation of FN-PSol neurons increased blood glucose levels. **b**, ITT after saline or CLZ injection (-15 min) into the mice (n = 5). **c**, Plasma C-peptide levels during GTT after saline or CLZ injection (-15 min) into the mice (n = 5). **d-g**, Oxygen consumption (VO₂, **d**), energy expenditure (**e**), carbohydrate utilization (**f**) and lipid utilization (**g**) measured in the calorimetry system. Mice were injected with saline or CLZ at 0 min (n = 10). Activation of FN-PSol neurons transiently reduced VO₂ (**d**) and energy expenditure (**e**). Activation of FN-PSol neurons decreased carbohydrate utilization (**f**) and increased lipid utilization (**g**). **h**, Open field test (OF) performed after saline or CLZ injection (-15 min) into the mice (n = 5). **i**, ITT (0–120 min) after CLZ injection (-15 min) of EYFP (n = 5) or hM4Di mice (n = 9), in which an inhibitory DREADD receptor was expressed in neurons projecting from the FN to the PSol. Data are presented as mean ± SEM; \**p* < 0.05, \*\**p* < 0.01, \*\*\**p* < 0.001, \*\*\*\**p* < 0.0001, two-way ANOVA followed by Sidak multiple comparison test in **a**, **e** and **g**; paired t-test in **h**.

**Extended Data Fig. 10.**
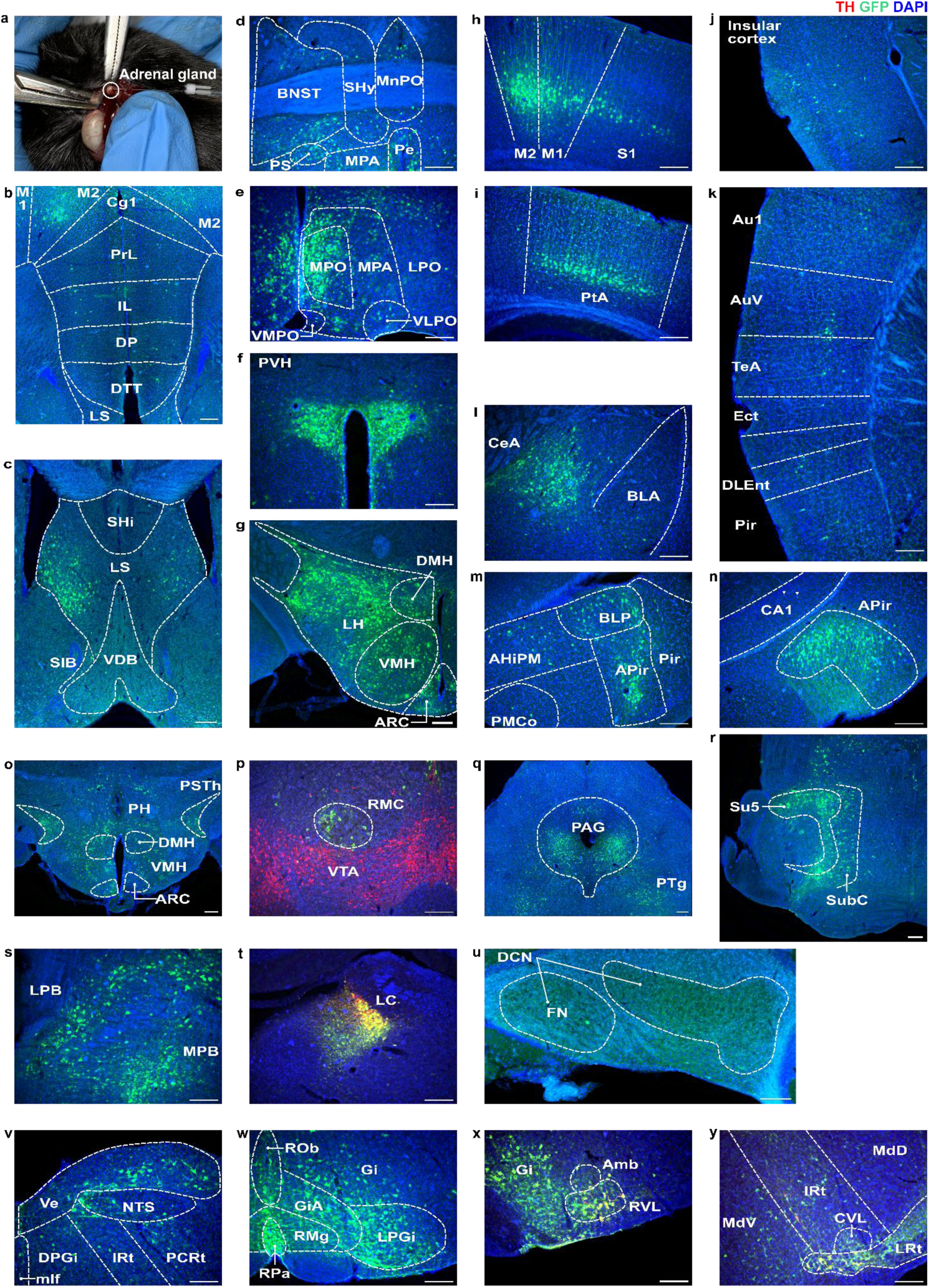
Neurons upstream of the adrenal gland. **a**, A picture of PRV injection into the adrenal gland. **b-y**, GFP expression in PRV infected cells in the brain. **p**, **t**, **x**, **y**, Tyrosine hydroxylase (TH) was stained as Red. **b**, M1, primary motor cortex; M2, secondary motor cortex; Cg1, cingulate cortex, area 1; PrL, prelimbic cortex; IL, infralimbic cortex; DP, dorsal peduncular cortex; DTT, dorsal tenia tecta; L, lateral septal nucleus. **c**, SHi, septohippocampal nucleus; VDB, the nucleus of the vertical limb of the diagonal band; SIB, substantia innominate, basal part. **d**, BNST, bed nucleus of the stria terminalis; SHy, septohypothalamic nucleus; MnPO, median preoptic nucleus; PS, parastrial; MPA, medial preoptic area; Pe, periventricular hypothalamic nucleus. **e**, MPO, medial preoptic nucleus; LPO, lateral preoptic area; VMPO, ventromedial preoptic nucleus; VLPO, ventrolateral preoptic nucleus. **f**, PVH, paraventricular hypothalamic nucleus. **g**, DMH, dorsomedial hypothalamic nucleus; VMH, ventromedial hypothalamic nucleus; ARC, arcuate hypothalamic nucleus; LH, lateral hypothalamus. **h**, S1, primary somatosensory cortex. **i**, PtA, parietal association cortex. **j**, Insular cortex. **k**, Au1, primary auditory cortex; AuV, secondary auditory cortex, ventral area; TeA, temporal association cortex; Ect, ectorhinal cortex; DLEnt, dorsolateral entorhinal cortex; Pir, piriform cortex. **l**, CeA, central amygdaloid nucleus; BLA, basolateral amygdaloid nucleus. **m**, AHiPM, amygdalohippocampal area, anterolateral part; BLP, basolateral amygdaloid nucleus, posterior part; APir, amygdalopiriform transition area. PMCo, posteromedial cortical amygdaloid area. **n**, CA1, field CA1 of the hippocampus. **o**, PH, posterior hypothalamic nucleus; PSTh, parasubthalamic nucleus. **p**, RMC, red nucleus, magnocellular part; VTA, ventral tegmental area, ventral tegmentum. **q**, PAG, periaqueductal gray; PTg, pedunculotegmental nucleus. **r**, Su5, supratrigeminal nucleus; SubC, subcoeruleus nucleus. **s**, LPB, lateral parabrachial nucleus; MPB, medial parabrachial nucleus. **t**, LC, locus coeruleus. **u**, DCN, deep cerebellar nucleus. **v**, Ve, vestibular nucleus; NTS, nucleus tractus solitarius; mlf, medial longitudinal fasciculus; DPGi, dorsal paragigantocellular nucleus; IRt, intermediate reticular nucleus; PCRt, parvicellular reticular nucleus. **w**, Rob, raphe obscurus nucleus; Gi, gigantocellular reticular nucleus; GiA, gigantocellular reticular nucleus, alpha part; LPGi, lateral gigantocellular reticular nucleus; RMg, raphe magnus nucleus; RPa, raphe palidus nucleus. **x**, Amb, ambiguous nucleus; RVL, rostral ventrolateral medulla. **y**, IRt, intermediate reticular nucleus; MdV, medullary reticular nucleus, ventral part; MdD, medullary reticular nucleus, dorsal part; CVL, caudoventrolateral reticular nucleus; LRt, lateral reticular nucleus. Scale bars are 200μm.

**Extended Data Fig. 11.**
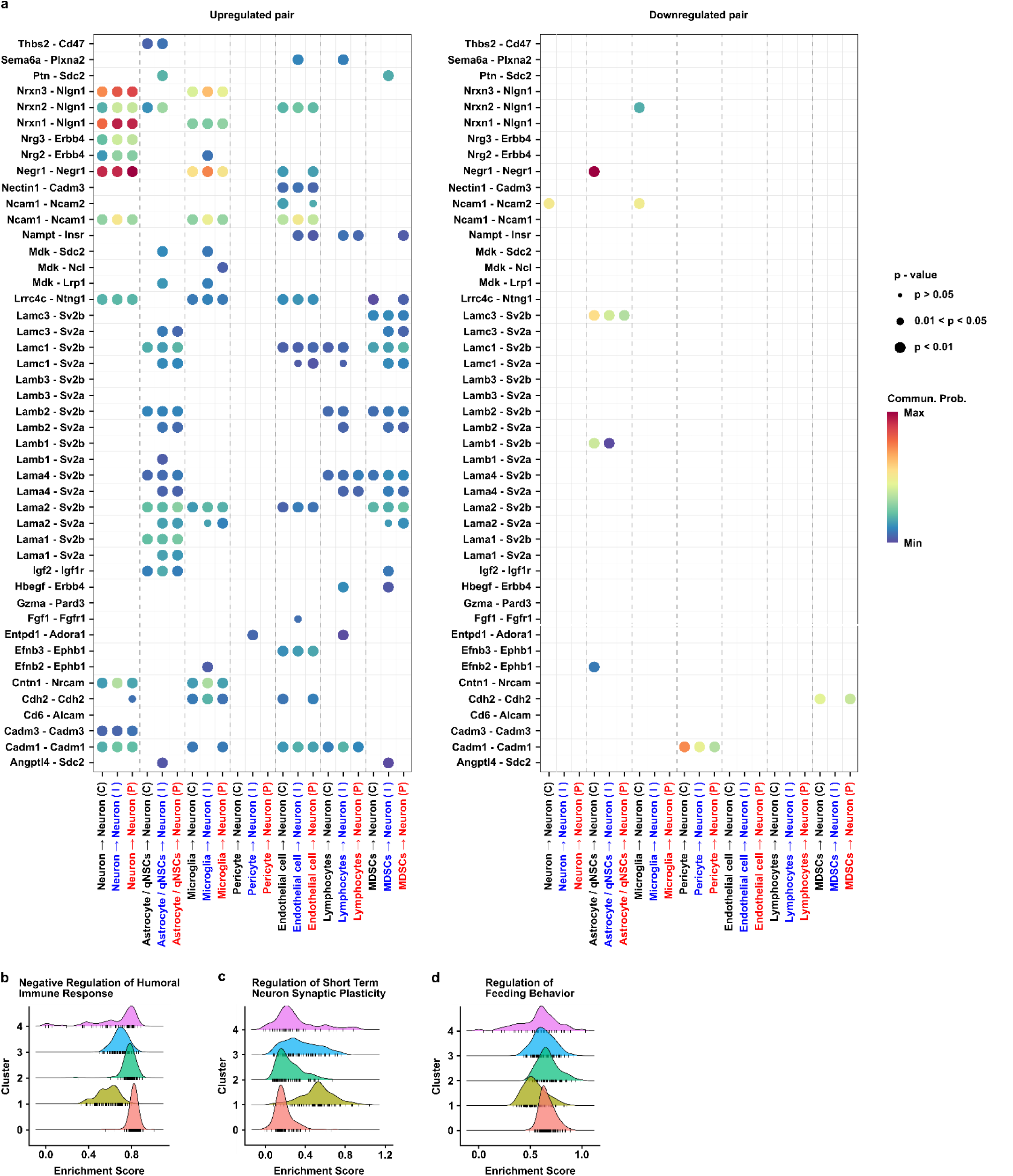
Analysis for cell-cell communication and GSEA. **a**, Cell-cell communication with neurons as receptors are shown for each cluster. Ligand-receptor pairs that were upregulated (left) or downregulated (right) by CSDS exposure are indicated. C, control; I, immediate-CSDS; P, post-CSDS. **b-d**, Gene set enrichment analysis (GSEA) showing GO terms with high enrichment scores in cluster 0 (**b**), cluster 1 (**c**), and cluster 2 (**d**).

**Supplementary Table 1.**
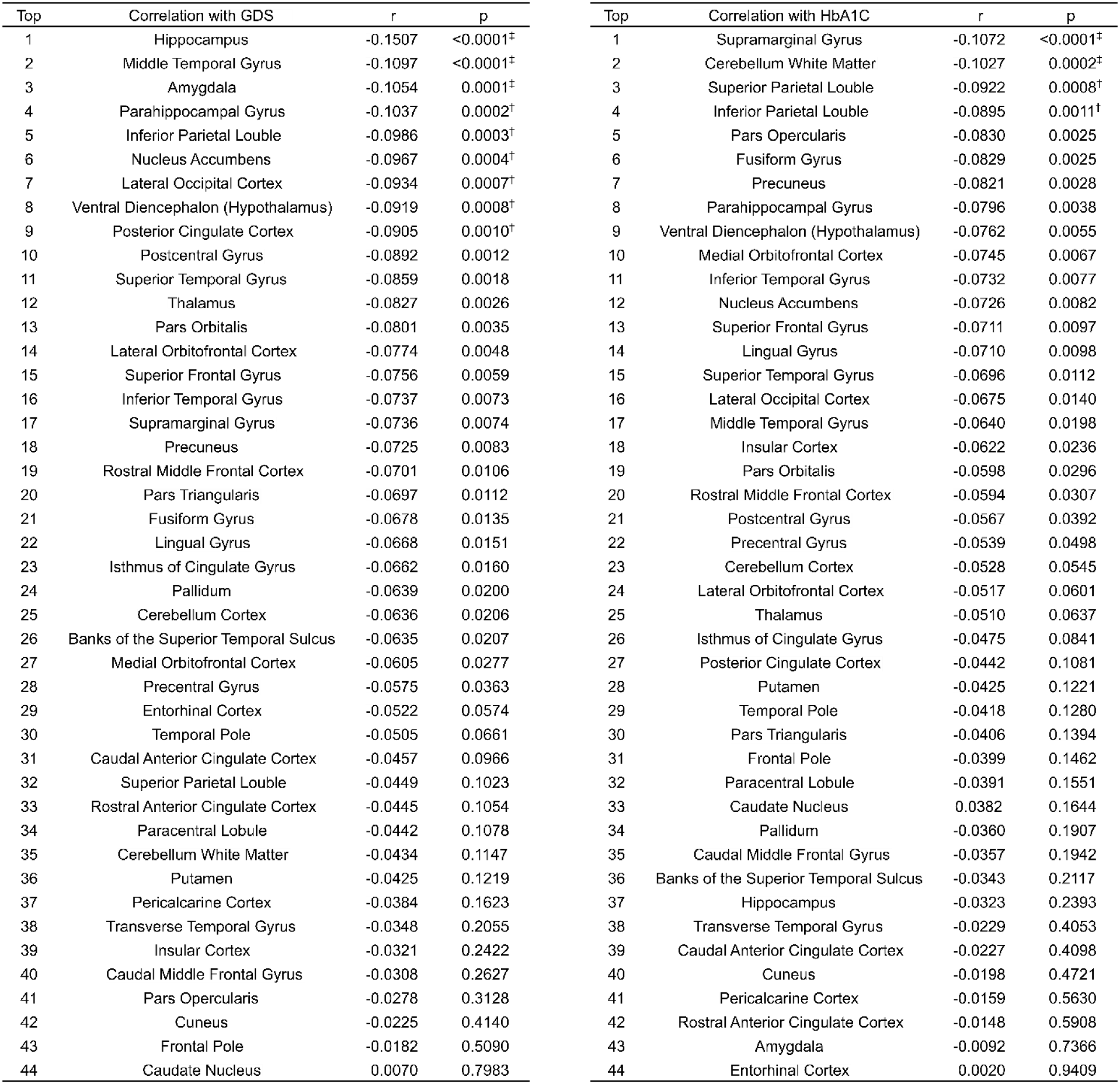
Correlation between brain volume and depression score or HbA1c. A study of the Arao cohort showed the correlation between brain region volumes and the Geriatric Depression Scale (GDS) or HbA1C. Each brain region’s volume was adjusted for estimated total intracranial volume. †*p* < 0.0011, ‡*p* < 0.0002, the p-values were calculated using Pearson correlation, and Bonferroni correction was applied to account for 44 multiple comparisons.

| Top | Correlation with GDS | r | p | Top | Correlation with HbA1C | r | p |
| --- | --- | --- | --- | --- | --- | --- | --- |
| 1 | Hippocampus | -0.1507 | <0.0001<sup>‡</sup> | 1 | Supramarginal Gyrus | -0.1072 | <0.0001<sup>‡</sup> |
| 2 | Middle Temporal Gyrus | -0.1097 | <0.0001<sup>‡</sup> | 2 | Cerebellum White Matter | -0.1027 | 0.0002<sup>‡</sup> |
| 3 | Amygdala | -0.1054 | 0.0001<sup>‡</sup> | 3 | Superior Parietal Lobule | -0.0922 | 0.0008<sup>†</sup> |
| 4 | Parahippocampal Gyrus | -0.1037 | 0.0002<sup>†</sup> | 4 | Inferior Parietal Lobule | -0.0895 | 0.0011<sup>†</sup> |
| 5 | Inferior Parietal Lobule | -0.0986 | 0.0003<sup>†</sup> | 5 | Pars Opercularis | -0.0830 | 0.0025 |
| 6 | Nucleus Accumbens | -0.0967 | 0.0004<sup>†</sup> | 6 | Fusiform Gyrus | -0.0829 | 0.0025 |
| 7 | Lateral Occipital Cortex | -0.0934 | 0.0007<sup>†</sup> | 7 | Precuneus | -0.0821 | 0.0028 |
| 8 | Ventral Diencephalon (Hypothalamus) | -0.0919 | 0.0008<sup>†</sup> | 8 | Parahippocampal Gyrus | -0.0796 | 0.0038 |
| 9 | Posterior Cingulate Cortex | -0.0905 | 0.0010<sup>†</sup> | 9 | Ventral Diencephalon (Hypothalamus) | -0.0762 | 0.0055 |
| 10 | Postcentral Gyrus | -0.0892 | 0.0012 | 10 | Medial Orbitofrontal Cortex | -0.0745 | 0.0067 |
| 11 | Superior Temporal Gyrus | -0.0859 | 0.0018 | 11 | Inferior Temporal Gyrus | -0.0732 | 0.0077 |
| 12 | Thalamus | -0.0827 | 0.0026 | 12 | Nucleus Accumbens | -0.0726 | 0.0082 |
| 13 | Pars Orbitalis | -0.0801 | 0.0035 | 13 | Superior Frontal Gyrus | -0.0711 | 0.0097 |
| 14 | Lateral Orbitofrontal Cortex | -0.0774 | 0.0048 | 14 | Lingual Gyrus | -0.0710 | 0.0098 |
| 15 | Superior Frontal Gyrus | -0.0756 | 0.0059 | 15 | Superior Temporal Gyrus | -0.0696 | 0.0112 |
| 16 | Inferior Temporal Gyrus | -0.0737 | 0.0073 | 16 | Lateral Occipital Cortex | -0.0675 | 0.0140 |
| 17 | Supramarginal Gyrus | -0.0736 | 0.0074 | 17 | Middle Temporal Gyrus | -0.0640 | 0.0198 |
| 18 | Precuneus | -0.0725 | 0.0083 | 18 | Insular Cortex | -0.0622 | 0.0236 |
| 19 | Rostral Middle Frontal Cortex | -0.0701 | 0.0106 | 19 | Pars Orbitalis | -0.0598 | 0.0296 |
| 20 | Pars Triangularis | -0.0697 | 0.0112 | 20 | Rostral Middle Frontal Cortex | -0.0594 | 0.0307 |
| 21 | Fusiform Gyrus | -0.0678 | 0.0135 | 21 | Postcentral Gyrus | -0.0567 | 0.0392 |
| 22 | Lingual Gyrus | -0.0668 | 0.0151 | 22 | Precentral Gyrus | -0.0539 | 0.0498 |
| 23 | Isthmus of Cingulate Gyrus | -0.0662 | 0.0160 | 23 | Cerebellum Cortex | -0.0528 | 0.0545 |
| 24 | Pallidum | -0.0639 | 0.0200 | 24 | Lateral Orbitofrontal Cortex | -0.0517 | 0.0601 |
| 25 | Cerebellum Cortex | -0.0636 | 0.0206 | 25 | Thalamus | -0.0510 | 0.0637 |
| 26 | Banks of the Superior Temporal Sulcus | -0.0635 | 0.0207 | 26 | Isthmus of Cingulate Gyrus | -0.0475 | 0.0841 |
| 27 | Medial Orbitofrontal Cortex | -0.0605 | 0.0277 | 27 | Posterior Cingulate Cortex | -0.0442 | 0.1081 |
| 28 | Precentral Gyrus | -0.0575 | 0.0363 | 28 | Putamen | -0.0425 | 0.1221 |
| 29 | Entorhinal Cortex | -0.0522 | 0.0574 | 29 | Temporal Pole | -0.0418 | 0.1280 |
| 30 | Temporal Pole | -0.0505 | 0.0661 | 30 | Pars Triangularis | -0.0406 | 0.1394 |
| 31 | Caudal Anterior Cingulate Cortex | -0.0457 | 0.0966 | 31 | Frontal Pole | -0.0399 | 0.1462 |
| 32 | Superior Parietal Lobule | -0.0449 | 0.1023 | 32 | Paracentral Lobule | -0.0391 | 0.1551 |
| 33 | Rostral Anterior Cingulate Cortex | -0.0445 | 0.1054 | 33 | Caudate Nucleus | 0.0382 | 0.1644 |
| 34 | Paracentral Lobule | -0.0442 | 0.1078 | 34 | Pallidum | -0.0360 | 0.1907 |
| 35 | Cerebellum White Matter | -0.0434 | 0.1147 | 35 | Caudal Middle Frontal Gyrus | -0.0357 | 0.1942 |
| 36 | Putamen | -0.0425 | 0.1219 | 36 | Banks of the Superior Temporal Sulcus | -0.0343 | 0.2117 |
| 37 | Pericalcarine Cortex | -0.0384 | 0.1623 | 37 | Hippocampus | -0.0323 | 0.2393 |
| 38 | Transverse Temporal Gyrus | -0.0348 | 0.2055 | 38 | Transverse Temporal Gyrus | -0.0229 | 0.4053 |
| 39 | Insular Cortex | -0.0321 | 0.2422 | 39 | Caudal Anterior Cingulate Cortex | -0.0227 | 0.4098 |
| 40 | Caudal Middle Frontal Gyrus | -0.0308 | 0.2627 | 40 | Cuneus | -0.0198 | 0.4721 |
| 41 | Pars Opercularis | -0.0278 | 0.3128 | 41 | Pericalcarine Cortex | -0.0159 | 0.5630 |
| 42 | Cuneus | -0.0225 | 0.4140 | 42 | Rostral Anterior Cingulate Cortex | -0.0148 | 0.5908 |
| 43 | Frontal Pole | -0.0182 | 0.5090 | 43 | Amygdala | -0.0092 | 0.7366 |
| 44 | Caudate Nucleus | 0.0070 | 0.7983 | 44 | Entorhinal Cortex | 0.0020 | 0.9409 |

**Supplementary Table 2.**
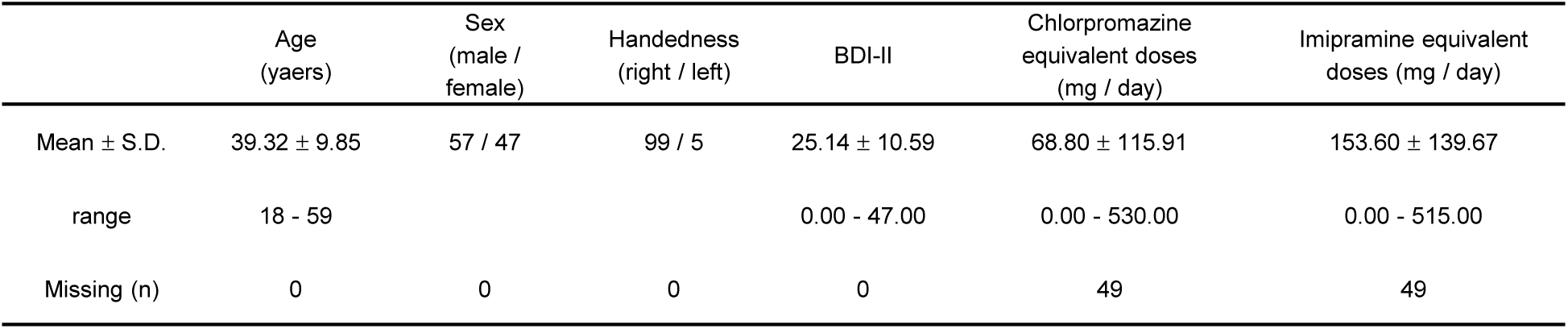
The characteristics of participants. Abbreviations: S.D., standard deviation, BDI-II, the Beck Depression Inventory-II.

**Supplementary Table 3.** Image acquisition parameters per procedure. AP, anterior-posterior; PA, posterior-anterior; TR, repetition time; TE, echo time COI, Siemens Verio scanner at the Center of Innovation in Hiroshima University; KUT, a Siemens TimTrio scanner at Kyoto University; UTO, GE MR750W scanner at The University of Tokyo Hospital.

|  | COI | Site<br>KUT | UTO |
| --- | --- | --- | --- |
| MRI scanner | Siemens verio | Siemens TimTrio | GE MR750w |
| Magnetic field strength | 3.0 T | 3.0 T | 3.0 T |
| Number of channels per coil | 12 | 32 | 24 |
| Field of view (mm) | 212 × 212 | 212 × 212 | 212 × 212 |
| Matrix | 64 × 64 | 64 × 64 | 64 × 64 |
| Phase encoding direction | AP | PA | PA |
| Number of slices | 40 | 40 | 40 |
| Slice thickness (mm) | 3.2 | 3.2 | 3.2 |
| Slice gap (mm) | 0.8 | 0.8 | 0.8 |
| TR (ms) | 2,500 | 2,500 | 2,500 |
| TE (ms) | 30 | 30 | 30 |
| Number of volumes | 240 | 240 | 240 |
| Total scan time (min:s) | 10:00 | 10:00 | 10:00 |
| Slice acquisition order | Ascending | Ascending | Ascending |
| n | 49 | 11 | 44 |

## Methods

### Animals

C57BL/6 and ICR male mice were purchased from SLC, Inc., Japan. Most experiments were conducted at the Graduate School of Veterinary Medicine, Hokkaido University. Studies involving the social interaction test and open field test were performed at the Graduate School of Medical Science, Kumamoto University. Some experiments involving inhibitory DREADD studies were conducted at the Graduate School of Pharmaceutical Sciences, Hokkaido University. Experiments involving indirect calorimetry were conducted at the Institute of Low-Temperature Science, Hokkaido University. The Pseudorabies virus studies were performed at the National Institute for Physiological Sciences and the National Institutes of Natural Sciences. Mice were maintained at 22– 24 °C and 30–60% humidity under a 12-h light/12-h dark cycle, except at the Institute of Low-Temperature Science, Hokkaido University, where they were housed under a 14-h light/10-h dark cycle. All mice had ad libitum access to food and water. Mice were fed laboratory chow, CE-2 (Oriental Yeast) at the Graduate School of Veterinary Medicine, Hokkaido University; Graduate School of Medical Science, Kumamoto University; and National Institute for Physiological Sciences, National Institutes of Natural Sciences; Labdiet 5053 (PMI, St. Louis) at the Graduate School of Pharmaceutical Sciences, Hokkaido University; MR stock (Nihon Nosan) at Institute of Low-Temperature Science, Hokkaido University. Animal care and experimental procedures were performed following guidelines and approval from the Animal Care and Use Committee of Hokkaido University, Kumamoto University, the University of Tokyo, or National Institute for Physiological Sciences, National Institutes of Natural Sciences.

### Chronic social defeat stress (CSDS)

We used ICR as resident mice and C57BL/6 as intruder mice based on previous reports^34^. The two were allowed to interact for 15 min, during which ICR attacked C57BL/6 (physical stress). After 15 min of physical stress, the ICR and C57BL/6 were separated using a partition with small holes (psychological stress). Stress exposure was repeated once a day for 10 days, using a different pair of ICR and C57BL/6 every day. On day 11, the C57BL/6 were housed individually.

### Social interaction test (SI)

We used a three-chambered social interaction test system (O’Hara & CO., LTD.). The field size was 40 cm × 61 cm and divided into three equally sized chambers by two partitions. A cage with a radius of 10 cm was placed in the corner of each side chamber. Mice could move freely between the chambers through an opening at the center of the partitions. On days 12 and 13 of the experiment, mice were allowed to freely explore the field for 5 min to habituate. During habituation, both side cages were empty. On day 13, the test mouse was moved to the test field, where one cage was placed with an ICR. The behavior was recorded for 5 min using a camera. The chamber containing the cage with the ICR was defined as the interaction zone, and the chamber on the opposite side was defined as the avoidance zone. The time spent in the avoidance or interaction chamber and the movement trajectory of the test mouse were analyzed using ImageJ and AnimalTracker^35^.

### Open Field test (OF)

We used a white box (38 cm × 38 cm) as the field. Mice were allowed to freely explore the field for 5 min, and their behavior was recorded with a camera (25 fps). In the CSDS study, the measurement was conducted once between days 11 and 16 of the experiment. In the chemogenetic study, saline or clozapine (CLZ, 0.1 mg/kg, Sigma-Aldrich) was administered intraperitoneally (i.p.) 15 min before the measurement. Experiments comparing saline with CLZ were conducted with an interval of more than three days. Mouse behavior was analyzed using ImageJ and MouBeAT^36^. The central 22.8 cm × 22.8 cm of the field was defined as the central zone. Immobility time was defined as the time during the experiment when the mouse’s movement was ≤ 0.1 cm/frame.

### Tolerance test

All tolerance tests were performed on ad libitum-fed mice after they moved to new cages. Blood glucose levels were measured with a glucose meter (Nipro). Mice were injected (i.p.) with glucose (2 g/kg), insulin (0.75 U/kg in Extended Fig. 9i, 0.5 U/kg in others), pyruvate (2 g/kg), glycerol (2.5 g/kg), or alanine solution (1 g/kg) at 0 min and measured blood glucose levels at 0, 15, 30, 60, and 120 min.

For chemogenetic studies, mice were injected (i.p.) with saline or CLZ (0.1 mg/kg) 15 min before tolerance tests. In Extended Fig. 5b, d, 9a, mice were injected with saline or CLZ at 0 min and measured blood glucose at 0, 15, 30, 45, 60, 75, and 120 min.

### Cannulation for Clamp Studies

For the hyperinsulinemic-euglycemic clamp (HE Clamp) and hyperglycemic clamp (HG Clamp) study, mice were anesthetized with a mixture of ketamine (100 mg/kg) and xylazine (10 mg/kg) and were cannulated in the right carotid artery and jugular vein one day before the measurement. These cannulas were routed subcutaneously to the dorsal side of the neck and the skin was closed^37^.

### Hyperinsulinemic-euglycemic clamp (HE Clamp)

The HE Clamp protocol was based on previous studies^38^. Mice were fed ad libitum, and the measurements were performed under freely moving conditions. During the Basal period (-90–0 min), [3-^3^H] glucose solution (0.05 μCi/min, Muromachi Kikai) was infused through a triple-lumen cannula connected to the jugular vein cannula. At the start of the Clamp period, in addition to [3-^3^H] glucose solution (0.1 μCi/min), an insulin solution (5.0 mU/kg/min, Novo Nordisk) was infused for 115 min. During the Clamp period, to maintain euglycemia (110–130 mg/dl), 30% glucose solution was infused through the triple-lumen cannula as required, and this was defined as the Glucose Infusion Rate (GIR). At 75 min, 2-[^14^C] Deoxy-D-glucose (2-[^14^C] DG, 10 μCi, Muromachi Kikai) was administered via the triple-lumen cannula. Blood was collected from the carotid artery cannula, and blood glucose levels were measured. Blood samples were collected at -15, -5, 5, 15, 25, 35, and 45 min. At the end of the experiment, the mice were euthanized via intravenous administration of thiopental (Nipro), and tissues including the prefrontal cortex, dorsal or ventral striatum, cortex, hypothalamus, hippocampus, amygdala, pons, brainstem, cerebellum, soleus, gastrocnemius (R, red muscle; W, white muscle), brown adipose tissue, heart, spleen, white adipose tissue, liver, and pancreas were rapidly collected and weighed.

The rate of disappearance (Rd), which reflects whole-body glucose utilization, was calculated from the plasma ^3^H-glucose (dpm/ml) concentration. Endogenous glucose production (EGP) was determined by subtracting GIR from Rd. Glycolysis was calculated using plasma ^3^H_2_O levels.

### Hyperglycemic clamp (HG Clamp)

During the HG Clamp, mice were fed ad libitum, and the measurements were performed under freely moving conditions. To maintain blood glucose levels at 250–300 mg/dl, a 30% glucose solution was infused as required through the jugular vein cannula for 120 min (0–120 min) as the GIR. Blood was collected from the carotid artery cannula, and blood glucose levels were measured at -15, -5, 5, 10, 20, 30, 40, 50, 60, 70, 80, 90, 100, 110, and 120 min, while blood samples were collected at -15, 20, 40, 60, 80, 100, and 120 min^37^.

### Measurements of blood hormones

The blood samples were centrifuged for 5 min at 2300 × *g* and maintained at −30°C until hormones were measured. Corticosterone ELISA kit (Enzo), Epinephrine/Norepinephrine ELISA kit (Abnova), and Mouse Insulin ELISA KIT (FUJIFILM Wako) were used. All the protocols followed the instructions provided by the kit.

### Indirect Calorimetry Analysis for Chemogenetic Study

Indirect calorimetry data for the chemogenetic study were measured using the ARCO-2000 system (ARCO system). Mice were acclimated to the measurement chambers one day prior to data collection. At 0 min, saline or CLZ (0.1 mg/kg) was injected (i.p.). On the following day, the mice were injected (i.p.) with the opposite solution (saline or CLZ) compared to the previous day. All values are presented as 10-minute averages.

### High-density multi-electrode arrays (HD-MEA)

We prepared aCSF by aerating 1× MaxOne solution with 95% O₂ and 5% CO₂, followed by adding CaCl₂ to achieve a concentration of 2 mM. Using vibratome, 300 µm sagittal cerebellum sections were corrected in ice-cold 1x MaxOne solution. After 30 min incubation in 37℃ aerated aCSF, 2-4 slices were recorded in each individual. Spikes were recorded at room temperature using the MaxOne HD-MEA system (MaxWell Biosystems AG) perfusing with aCSF aerated with 95% O2 and 5% CO_2_. Recording electrodes were selected based on the results of an activity scan. Subsequently, spikes in the aCSF solutions with 7.25mM K^+^, GABA antagonist (10µM bicuculline), glutamate antagonist (10µM CNQX, 50µM DL-AP5), were recorded. Spike sorting was performed using UMAP dimensionality reduction with graph clustering in MATLAB (version R2019b, MathWorks, Natick), excluding data sets with fewer than 100 spikes. The inter-spike interval (ISI) was calculated for each isolated single-unit, and the mode of the ISI distribution was compared between control and post-CSDS mice across different solutions.

### In vivo extracellular recordings of FN firing in unanesthetized mice

Each mouse was anesthetized with 1–2% isoflurane and placed in a conventional stereotaxic apparatus. Under anesthesia, the skull was exposed, and a U-shaped head holder was fixed on the skull with bone-adhesive resin^39^. After recovery, each mouse was habituated to the stereotaxic apparatus through repeated head fixation sessions. On the recording day, each mouse was initially anesthetized and mounted on a stereotaxic apparatus with its head fixed. A cranial window was made, and a silicon probe (A1×32-Poly2-5mm-50s-177, NeuroNexus) was perpendicularly inserted above the target region (AP: -6.24 mm, ML: 0.75 mm from the bregma, DV: 2.0 mm from the dura), and then gently advanced into the FN (DV: 2.53–2.97 mm from the dura). Once stable recordings were observed, anesthesia was discontinued. After a 30-minute recovery period, data acquisition recordings were initiated. Neural signals were amplified using a RHD recording headstage (#C3314, Intan Technologies) and recorded at 30 kHz via the Open Ephys acquisition system. Neuronal spikes were detected from high-pass filtered raw signals (0.5–5 kHz) by a threshold crossing-based algorithm. Detected spikes were automatically sorted using klusta software^44^. This automatic clustering process was followed by the manual refinement of the clusters using phy software (https://github.com/cortex-lab/phy).

### Viruses

All AAVs were obtained from addgene or UNC and diluted with PBS. AAV and final concentration were AAV-hSyn-hM3Dq-mCherry (addgene, 2.4 × 10^12^ GC/ml), AAV2-hSyn-hM4DGi (addgene, 2.0 × 10^12^ GC/ml), AAV2-hSyn-mCherry-Cre (addgene, 2.0 × 10^12^ GC/ml or UNC, 5.0 × 10^12^ GC/ml), AAV2-Ef1a-DIO-hChR2 (H134R)-EYFP (UNC, 1.8 × 10^12^ GC/ml), AAV2-hSyn-DIO-hM3Dq-mCherry (addgene, 4.5 × 10^12^ GC/ml), AAV2-hSyn-DIO-hM4Di-mCherry (addgene, 2.8 × 10^12^ GC/ml), AAVrg-Ef1a-mCherry-IRES-Cre (addgene, 3.9 × 10^12^ GC/ml), AAVrg-CAG-hChR2 (H134R)-tdTomato (addgene, 4.5 × 10^12^ GC/ml). To obtain the PRV expressing GFP (PRV152), BHK21 cells (Japanese Collection of Research Bioresources Cell Bank (JCRB), #JCRB9020) were infected with the parental viruses (kindly provided by Lynn Enquist (Princeton University)) with a multiplicity of infection (M.O.I.) = 0.1∼0.01. Once a prominent cytopathic effect was observed in infected cells, the cell media was harvested, centrifuged at 1000 × *g* for 5 minutes to remove cell debris, and subjected to the ultracentrifugation with 30% sucrose cushion at 68,000 × *g* for 2 hours to concentrate the virus. The viral pellet was then resuspended in HBSS. Viral titers were determined using standard plaque assays with PK15 cells (ATCC, #CCL-33).

### Stereotaxic surgeries and AAV injection

Male C57BL/6 mice (8–9 weeks old) were anesthetized with a mixture of ketamine (100 mg/kg) and xylazine (10 mg/kg) at the Graduate School of Veterinary Medicine, Hokkaido University, with isoflurane (0.8–1.5%) at the Graduate School of Pharmaceutical Sciences, Hokkaido University; with three types of mixed anesthetic agents at the Institute of Low-Temperature Science, Hokkaido University. For chemogenetic studies focused on all FN neurons, mice were injected with AAV2-hSyn-hM3Dq-mCherry or AAV2-hSyn-hM4DGi or a mixture of AAV2-hSyn-mCherry-Cre and AAV2-Ef1a-DIO-hChR2 (H134R)-EYFP into the FN (AP: -6.24 mm, L: ± 0.75 mm, DV: 3.50 mm). For chemogenetic studies focused on FN neurons that innerve specific brain regions, mice were injected with AAV2-hSyn-DIO hM3Dq-mCherry or AAV2-hSyn-DIO-hM4Di-mCherry or AAV2-Ef1a-DIO-hChR2 (H134R)-EYFP in FN and injected AAVrg-Ef1a-mCherry-IRES-Cre in PSol (AP: -7.32 mm, L: ± 0.70 mm, DV: 4.30 mm), POA (AP: 0.38 mm, L: ± 0.30 mm, DV: 5.25 mm), PVT (AP: -1.06 mm, L: ± 0.00 mm, DV: 3.10 mm), LH (AP: -1.46 mm, L: ± 1.10 mm, DV: 5.3 mm), DMH (AP: -1.70 mm, L: ± 1.10 mm, DV: 5.30 mm), PF (AP: -2.10 mm, L: ± 0.70 mm, DV: 3.42-3.00 mm), PB (AP: -5.4 mm, L: ± 1.60 mm, DV: 3.55 mm). For the anatomical study in Fig. 2t, mice were injected with a mixture of AAV2-hSyn-mCherry-Cre and AAV2-Ef1a-DIO-hChR2 (H134R)-EYFP into the FN and injected with AAVrg-CAG-hChR2 (H134R)-tdTomato into the RVL (AP: -7.10 mm, L: ± 1.20 mm, DV: 5.90 mm). All AAVs were injected into both sides of the brain regions. The volume of AAV on one side was 0.6 µl in the LH and 0.3 µl in the others. Brains were collected to check the injection site. The mice, in which the AAV injection was not successful were removed from the data. The waiting period for recovery and virus expression for the experiments was at least 4 weeks.

### DREADD agonist

CLZ was used as the DREADD agonist. A dose of 0.1 mg/kg was selected to ensure sufficient DREADD activation while avoiding the side effects of hyperglycemia^40,41^.

### PRV injection

Male C57BL/6 mice were anesthetized with intraperitoneal injection of a mixture of ketamine (100 mg/kg) and xylazine (10 mg/kg). Mice were placed in a prone position to access the left adrenal gland and injected with 0.3 µl PRV (1.5 × 10^10^ pfu/m1) into the left adrenal gland. The skin incision was sutured after injection. Mice were monitored daily and euthanized for 5 days after surgery.

### Sectioning and Immunohistochemistry

Mice were euthanized using CO_2_ or isoflurane and perfused transcardially with heparinized saline followed by 4% paraformaldehyde (PFA). Brains were collected and subsequently immersed in 4% PFA for one day, followed by 30% sucrose solution in 0.1 M phosphate buffer (PB) for another day. Brain sections were sliced at a thickness of 50 μm using a cryostat (Leica). For immunohistochemical staining to enhance EYFP or GFP signals, floating sections were incubated for 1 h in a blocking solution (4% normal goat serum, 0.4% Triton-X100, 1% bovine serum albumin, and 0.1% glycine in 0.1 M PB). After washing, sections were incubated overnight in Goat-anti-GFP antibody (1:1000, ROCKLAND) in 0.1 M PB. The sections were washed and incubated for 2 h at room temperature with Alexa 488 Donkey Anti-Goat (IgG) secondary antibody (1:500, ab150129, Abcam). In immunohistochemical staining for TH, Rabbit-anti-Tyrosine Hydroxylase antibody (1:1000, AB152, Merck Millipore) was used for the first antibody, Anti-rabbit IgG (H+L), F(ab’)_2_ Fragment (1:500, Alexa Fluor 594, Cell Signaling Technology) was used as the second antibody. The protocol was the same as that for GFP staining. The sections were mounted using a nuclear staining mounting medium (DAPI Fluoromount-G, Southern Biotechnology). Images were acquired using an all-in-one fluorescence microscope (BZ-9000 or BZ-X710, Keyence).

### Sample collection for scRNAseq

In the scRNAseq study, transcriptome analysis was conducted by pooling DCN samples from control (n = 8), immediate-CSDS (n = 8), and post-CSDS (n = 8) mice into one tube per group. Mice were euthanized using CO_2_, and their brains were collected. The brains were sectioned in aCSF (124 mM NaCl, 3.0 mM KCl, 2.0 mM CaCl_2_-2H_2_O, 2.0 mM MgCl_2_-6H_2_O, 1.23 mM NaH_2_PO_4_-H_2_O, 26 mM NaHCO_3_, 10 mM Glucose) that had been aerated with O_2_ for over an hour, and the DCN was isolated. The DCN tissue was shaken in trituration solution (aCSF with 0.3 U/ml Papain, 0.075 µg/ml, 3.75 µg/ml BSA) and centrifuged. The tissue was washed in aCSF containing TTX (100 nM), DQNX (20 µM), APV (50 µM), and 10% FBS and triturated. The tissue was passed through a 40 µm mesh filter, washed again with aCSF containing TTX, DQNX, APV, and 10% FBS, and adjusted to a total volume of 4 ml. Calcein-AM (250 nM, Sigma-Aldrich) was added, and live cells were sorted using a cell sorter. The scRNAseq libraries were prepared using Chromium NextGEM Single Cell 3’ Gel Bead Kit v3.1 (10x Genomics). All libraries were sequenced on MGI DNBSEQ-G400 platform with 2 × 100 bp paired end mode.

### Processing of scRNA-seq data

Raw reads were aligned to the GRCm38 reference genome, UMI (unique molecular identifier) counting was performed using Cell Ranger (version 7.0.1). Seurat (version 4.3.0.1) was used for quality filtering and downstream analysis. Low-quality cells (≤ 300 genes/cell, ≥ 10% mitochondrial genes/cell) were excluded. Potential doublets were removed by DoubletFinder (version 2.0.3). To integrate “control”, “immediate-CSDS” and “post-CSDS” samples, we used Seurat’s anchoring integration method. We performed principal component analysis (PCA) and graph-based Louvain clustering on the top 20 principal components (PCs). The cluster-specific marker genes were identified using FindAllMarkers function. The cell clusters were manually annotated according to these marker genes. Erythrocytes were removed from the data. Clustering results were visualized on uniform manifold approximation and projection (UMAP) plots.

### Cell-cell interaction analysis

Cell-cell interactions were inferred using the CellChat (version 1.6.1). The netVisual_bubble function was used to visualize the CSDS-upregulated interactions in ligands (originating from any cell types) and receptors (in the neuron subsets). The netVisual_individual function was used to visualize the individual ligand-receptor pair which showed significant interaction (P-value < 0.05).

### Gene set enrichment analysis (GSIS)

To functionally describe neuron cell subtypes, we performed GSIS on the scRNA-seq dataset using the “ssGSEA” method from the escape (version 2.0.0) and Gene Ontology Biological Processes term.

### The Arao cohort

We analyzed data from the Arao cohort, a subset of the Japan Prospective Studies Collaboration for Aging and Dementia (JPSC-AD)^42^. This cohort includes residents aged ≥65 years (mean ± SD, 73.75 ± 6.21) surveyed in Arao City, Kumamoto Prefecture. Baseline data from 1,325 participants (Male, n = 505; Female, n = 820) with complete records for the Geriatric Depression Scale (GDS), HbA1C, and structural MRI were used, after excluding those with missing data, traumatic brain injury, dementia, or stroke. MRI measurements were conducted at the Arao Municipal Hospital (Kumamoto, Japan, n = 877) and Omuta Tenryo Hospital (Fukuoka, Japan, n = 448). Brain region volumes were calculated using FreeSurfer version 5.3 with the Desikan-Killany Atlas and normalized by eTIV (estimated total intracranial volume). Participants were categorized into three groups based on GDS scores: normal (0–4, n = 1,140), mild depression (5–9, n = 165), and depression (10–15, n = 20). The study was approved by the ethics committee of Kumamoto University (GENOME-333), and written informed consent was obtained.

### fMRI Study on patients with depression

#### Participants

Total 104 participants clinically diagnosed with major depressive disorder (MDD) were collected from the database of the Japanese Strategic Research Program for the Promotion of Brain Science (SRPBS) Decoded Neurofeedback (DecNef) Consortium^43,44^, and additional brain images scanned in the Department of Psychiatry, The University of Tokyo (Supplementary Table 2). The detailed inclusion and exclusion criteria have been previously described^43^. This study was approved by the appropriate institutional review boards^43^. All participants provided written informed consent. The severity of depressive symptoms was assessed using the Japanese version of the Beck Depression Inventory-II (BDI-II)^45,46^.

#### Resting-state functional magnetic resonance imaging data acquisition

Resting-state functional magnetic resonance imaging (rs-fMRI) data were acquired using three scanners (Supplementary Table 3). We instructed the participants to relax but not to sleep during scanning and to focus on the central crosshair mark.

#### Image preprocessing

Image preprocessing was performed using Statistical Parametric Mapping (SPM12, v7771; Wellcome Department of Cognitive Neurology) in Matlab R2019b (Mathworks, Natick). Conventional preprocessing was performed. First, slice timing correction and geometric distortion correction^47^ were conducted for functional images. Then, the participant’s high-resolution T1-weighted anatomical image was coregistered to their functional images. The coregistered anatomical image was processed using a unified segmentation procedure combining segmentation, bias correction, and spatial normalization into a standard template (Montreal Neurological Institute). The same normalization parameters were used to normalize the functional images. We excluded participants with an estimated head-motion exceeding 3 mm in any direction from the analysis. This is because 1 voxel size was approximately 3 × 3 × 3 mm^3^.

Furthermore, several of our regions of interest were separated by small distances (in the order of a few millimeters). This called for a spatially precise region of interest definition that was not confounded by head movement. Normalized functional images were smoothed in space with a 6-mm full-width at half-maximum 3D isotropic Gaussian kernel and high-pass filtered with a 128 s (0.01 Hz) cut-off to remove low-frequency drifts. Furthermore, we calculated the derivative or root mean square variance over voxels (DVARS)^48^, quantifying the mean change in image intensity between the time points. We used the DVARS and six rigid motion parameters for the preprocessed fMRI time series to regress out the effects of head motion. Subsequently, time series were extracted from the white matter and cerebrospinal fluid, and those time series were regressed out from preprocessed fMRI data to control for the effect of physiological noise.

#### Region of interest

The fastigial nuclei mask was created using the cerebellar atlas^51^ of the JuBrain Anatomy Toolbox^49–51^. Then, a 6 mm-radius sphere centered on the center of the fastigial nuclei mask ([x, y, z] = [0, -54, -30]) was created as a region of interest (ROI).

#### Seed-based connectivity maps and associations with depressive symptoms

At the individual level, first, time series were extracted from the fastigial nuclei ROI for individual preprocessed rs-fMRI data. Then, the general liner model (GLM) was created using the extracted time series as a statistical regressor against whole-brain rs-fMRI data to identify brain voxels that showed a significant correlation with the extracted time series data from the ROI (seed-based connectivity maps). At the group level, the associations between individual seed-based connectivity maps and BDI-II values were examined using the GLM using BDI-II values as a statistical regressor against seed-based connectivity maps. As rs-fMRI data were obtained using three different scanners, the effect of the scanner was included in the GLM as a confounder. As there was no significant effect of sex on BDI-II and no significant correlation between age and BDI-II, sex and age were not included in the GLM as confounders. For the group level analysis, a cluster defining threshold (CDT) of z = 3.1 was used to determine whether a cluster of voxels was significant. Simulations were run to see how often we would get clusters of a certain size with each of their constituent voxels passing this z-threshold, and a distribution of cluster sizes is generated for that CDT. Cluster sizes that occur less than 5% of the time in the simulations for that CDT are then determined to be significant.

#### Statistical analysis and reproducibility

Sample sizes are provided in the figure legends. Measurements are expressed as mean ± SEM. A two-tailed t-test was used for comparisons between two independent groups. A paired t-test was used for paired data comparisons in excitatory DREADD studies, in which the same individuals were used. Pearson correlation was used to assess linear correlations. One-way ANOVA followed by Sidak’s multiple comparison test was used for comparisons across three groups. One-way ANOVA followed by Tukey’s multiple comparison test, as used in Extended Data Fig. 4c, was performed specifically for comparisons among samples where only one experimental condition differed between the groups. Two-way ANOVA followed by Sidak’s multiple comparison test was used to analyze temporal changes between two groups. These analyses were performed using GraphPad Prism 10 (GraphPad Software, Inc.). Two-sample Kolmogorov-Smirnov test was performed in MATLAB. The generalized linear mixed model was analyzed using a log-normal distribution in JMP Pro 18.

## Data availability

The data that supports the findings of this study are available from the corresponding author upon reasonable request. For the rs-fMRI dataset are provided from the SRPBS Multidisorder Dataset and please request via https://bicr.atr.jp/decnefpro/data/.

## Author contributions

C.T. conceived this study, designed the experiments, and supervised the entire study. T.I. performed most of the experiments and analysis. T.A., K.S. contributed HD-MEA. Y.N., S.K., K.K., performed fMRI study. T.T., K.Y. analyzed scRNA seq. K.K. contributed PRV study. N.K., M.T. performed Arao cohort. K.Y., Y.T., M.M., contributed in vivo extracellular recording. K.X.K, S.X., M.K., M.I., T.I. contributed behavior tests.Y.Y. contributed indirect calorimetry. T.I, C.T. wrote the manuscript. J.J.Y. assisted in preparing the manuscript.

